# A cell non-autonomous FOXO/DAF-16-mediated germline quality assurance program that responds to somatic DNA damage

**DOI:** 10.1101/2022.03.03.482777

**Authors:** Gautam Chandra Sarkar, Umanshi Rautela, Anita Goyala, Sudeshna Datta, Nikhita Anand, Anupama Singh, Prachi Singh, Manish Chamoli, Arnab Mukhopadhyay

## Abstract

Germline integrity is critical for progeny fitness. Organisms deploy the DNA damage response (DDR) signalling to protect germline from genotoxic stress, facilitating cell-cycle arrest of germ cells and DNA repair or their apoptosis. Cell-autonomous regulation of germline quality is well-studied; however, how quality is enforced cell non-autonomously on sensing somatic DNA damage is less known. Using *Caenorhabditis elegans*, we show that DDR disruption, only in the uterus, when insulin-IGF-1 signalling (IIS) is low, arrests germline development and induces sterility in a FOXO/DAF-16 transcription factor (TF)-dependent manner. Without FOXO/DAF-16, germ cells of the IIS mutant escape arrest to produce poor quality oocytes, showing that the TF imposes strict quality control during low IIS. In response to low IIS in neurons, FOXO/DAF-16 works cell autonomously as well as non-autonomously to facilitate the arrest. Activated FOXO/DAF-16 promotes transcription of checkpoint and DDR genes, protecting germline integrity. However, on reducing DDR during low IIS, the TF decreases ERK/MPK-1 signaling below a threshold, and transcriptionally downregulates genes involved in spermatogenesis-to-oogenesis switch as well as *cdk-1*/Cyclin B to promote germline arrest. Altogether, our study reveals how cell non-autonomous function of FOXO/DAF-16 promotes germline quality and progeny fitness in response to somatic DNA damage.

**Significance Statement:** Reproductive decisions are supervised processes that take into account various inputs like cellular energy availability and status of damage repair in order to ensure healthy progeny. In this study, we show that the absence of optimal DNA damage repair in the somatic uterine tissues prevents oocyte development by the cell-autonomous as well non-autonomous function of activated FOXO transcription factor DAF-16. Thus, this study elucidates a new surveillance role of FOXO/DAF-16 in somatic tissues that ensures progeny fitness.

## Introduction

The propagation of a species depends on a healthy and productive germline. The stability of the genome is constantly under threat from extrinsic as well as cell-intrinsic genotoxic agents. Thus, all organisms invest heavily on protecting the germline against DNA damage. Generally, in response to DNA damage, an organism deploys an array of countermeasures. Depending on the type of DNA damage, organisms employ lesion-specific DNA repair pathways that can restore damage inflicted by ultra-violet rays (UV), ionizing radiation (IR) or reactive oxygen (ROS) and nitrogen species (RNS). Apart from these highly specialized DNA repair mechanisms, organisms also depend on DNA damage response (DDR) signaling to activate damage-responsive checkpoints, leading to cell cycle arrest to repair the damage or apoptosis, when damage is beyond repair. Perturbation of the DDR, in turn, leads to unrepaired DNA damage, genomic instability and are the basis of many human diseases like cancer, neurodegeneration as well as aging (1). Unrepaired DNA lesions in the germline can lead to infertility, reduced progeny fitness and birth defects. The critical decision of reproductive commitment and germline proliferation is largely influenced by environmental conditions via soma to germline communication (2, 3). For example, irradiation (genotoxic stress) of somatic tissues has been shown to cause hormonal imbalance leading to increased incidences of infertility in female cancer patients(4). However, it is less known whether or how an organism perceives intrinsic DNA damage signals in somatic tissues and regulates germline development to preserve progeny genome integrity

Research in *C. elegans* has elucidated the role of the conserved FOXO TF DAF-16 in somatic and germline quality assurance. Mutations in the neuroendocrine IIS pathway activate FOXO/DAF-16 to arrest development at dauer diapause (5, 6). The TF mediates arrest at the L1 larval stage when food is depleted (7). Further, activated FOXO/DAF-16 delays aging, enhances resistance to stresses and increases life span under conditions of lowered IIS (8), (9, 10). These IIS mutant animals maintain their germline stem cell pool even at an advanced age, and so, have delayed reproductive aging (11). They produce better quality oocytes (12) with low chromosomal abnormalities as compared to wild-type (WT), but the mechanism is less understood (13). Interestingly, the IIS receptor DAF-2 functions cell non-autonomously in the neuron whereas DAF-16 works in the intestine to regulate longevity (14, 15). The long reproductive span or higher oocyte quality of the *daf-2* mutant is dependent on muscle or intestinal DAF-16 activity (13). However, it is not known whether activated FOXO/DAF-16 can sense DNA damage in somatic tissues and modulate germline development cell non-autonomously.

Here, we show that in *C. elegans*, a uterine tissue-specific perturbation of DDR in the IIS pathway mutants prevents germ cells from exiting the pachytene stage of meiosis and inhibits oogenesis. For disruption of DDR and inducing DNA damage, we knocked-down (KD) *cdk-12* that is required for the transcription of DDR genes (16, 17). This sterility is reversed in the absence of DAF-16, leading to the production of poor-quality oocytes and developmentally retarded progeny. We elucidate the cell autonomous as well as non-autonomous requirements of the IIS pathway and FOXO/DAF-16 in orchestrating the arrest. We show that this is achieved by downregulating signaling of ERK-MPK-1 pathway along with the transcriptional downregulation of important genes required for germline development. Thus, our study elucidates a new cell non-autonomous role of the IIS pathway and FOXO/DAF-16 in ensuring germline quality in response to somatic perturbation of DDR and associated chance of genome instability in the progeny.

## Results

### The cyclin-dependent kinase gene *cdk-12* genetically interacts with the IIS pathway

We were interested in identifying genes that when knocked down induce chronic stress signaling, thereby enhancing dauer formation of the IIS receptor mutant *daf-2(e1370)* (referred to as *daf-2*) strain. Knocking down *cdk-12* using RNAi led to a significant increase in dauer formation (**Figure 1A**). In line with its possible role in inducing stress, *cdk-12* RNAi, initiated at L4, reduced life span of wild-type (WT), *daf-2*, *daf-16(mgdf50)* (referred to as *daf-16*) and *daf-16;daf-2* to an equal extent **(Figure S1A-D, Table S1)**. Thus, CDK-12 depletion may cause chronic stress to the worms, thereby increasing dauer of *daf-2* and reducing life span in general.

**Figure 1.**
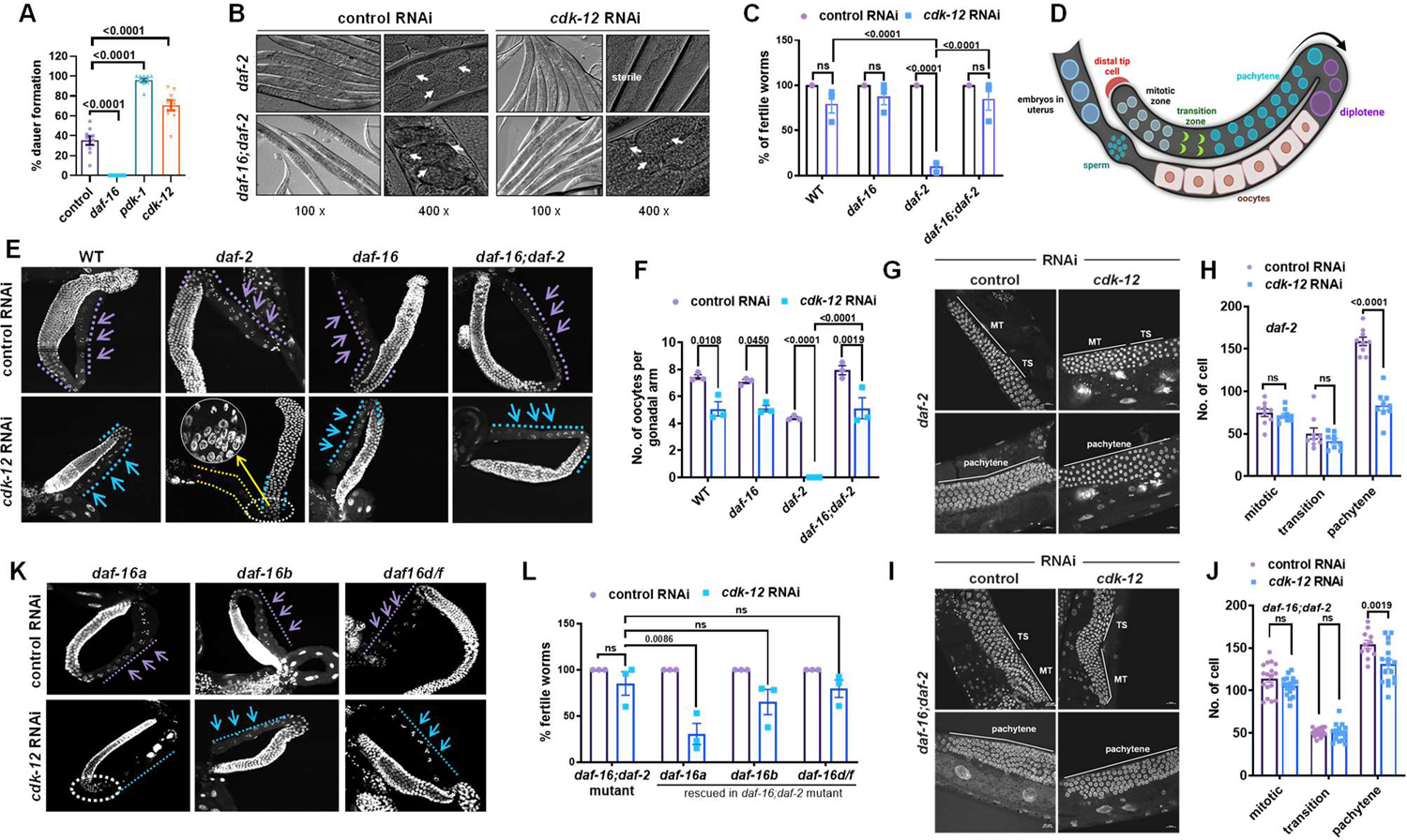
CDK-12 KD arrests germline of IIS mutant in a FOXO/DAF-16-dependent manner. (A) Percentage of dauer formation in *daf-2(e1370)* when *daf-16*, *pdk-1* or *cdk-12* is knocked down (KD) using RNAi. *Cdk-12* KD increased dauer formation of *daf-2(e1370).* Average of nine biological replicates (n≥40 for each replicate). One way ANOVA. Each point represents mean percentage of dauer formation for one biological replicate. Experiments performed at 22.5°C (B) Representative images showing that *cdk-12* RNAi results in sterility in *daf-2(e1370)* worms that is rescued in *daf-2(e1370)*;*daf-16(mgdf50)*. Arrows show eggs. Image were captured at 100X and 400X magnification for each condition. (C) Percentage of fertile worms in wild-type (WT), *daf-16(mgdf50), daf-2(e1370)* and *daf-16(mgdf50);daf-2(e1370)* on *cdk-12* RNAi. Most of the *daf-2(e1370)* worms are sterile on *cdk-12* KD that is rescued in *daf-16(mgdf50);daf-2(e1370)*. Average of three biological replicates (n≥25 for each experiment). Two-way ANOVA-Sidak multiple comparisons test. (D) A diagrammatic representation of one of the two arms of the *C. elegans* gonad. (E) Representative fluorescence images of dissected gonadal arms that were stained with DAPI. The germline arrests at the pachytene stage of meiosis 1 in *daf-2(e1370)* worms upon *cdk-12* KD; this was rescued in *daf-16(mgdf50);daf-2(e1370).* Image were captured at 400X magnification. (F) Oocyte counts in WT, *daf-16(mgdf50), daf-2(e1370)* and *daf-16(mgdf50);daf-2(e1370)* on *cdk-12* RNAi. Average of three biological replicates (n≥15 for each experiment). Two-way ANOVA-Sidak multiple comparisons test. (G-J) Representative fluorescence images of DAPI-stained dissected gonads of *daf-2(e1370)* and *daf-16(mgdf50);daf-2(e1370)* on control or *cdk-12* RNAi, showing germ cells in mitotic, transition and pachytene zones (G, I) and their quantification (H, J). n=9 (*daf-2*), n=17 (*daf-16;daf-2*) gonads for each condition used in quantification. One way ANOVA. Each point represents the number of mitotic (MT), transition (TS) or pachytene zones cell. (K) Representative fluorescence images of DAPI-stained dissected gonads of *daf-16(mgdf50);daf-2(e1370)* worms, which have been transgenically rescued with different *daf-16* isoforms *(daf-16a, daf-16b* or *daf-16d/f)*, when grown on control or *cdk-12* RNAi. Arrows showing the oocytes. Image were captured at 400X magnification. (L) Percentage of fertile worms in *daf-16(mgdf50);daf-2(e1370)* that are rescued with *daf-16* isoforms *(daf-16a, daf-16b* or *daf-16d/f)* on control or *cdk-12* RNAi. Average of three biological replicates (n≥40 for each replicate). Two-way ANOVA-Sidak multiple comparisons test. Error bars are SEM. ns, non-significant. Unless otherwise mentioned, all experiments were performed at 20 °C. Source data is provided in Dataset S1.

### *Cdk-12* depletion during low IIS leads to DAF-16A isoform-dependent pachytene arrest of germline and sterility

Considering *cdk-12* knockdown may potentially induce stress, we asked whether this would affect progeny production. Interestingly, we found that the *daf-2* worms became sterile when they were grown on *cdk-12* RNAi from L1 onwards (**Figure 1B, C**). The sterility is DAF-16-dependent as fertility was restored in *daf-16;daf-2*, signifying that DAF-16 regulates the germline arrest in *daf-2* (**Figure 1B, C**). Importantly, this was not due to differential RNAi efficiency in the strains **(Figure S1E).**

The *C. elegans* hermaphrodite gonad has two U-shaped arms carrying germline stem cell (GSC) pool near the distal end, which divide mitotically and then enter meiotic prophase as they move away from the distal tip. Germ cells in meiosis produce sperms during larval 4 (L4) stage, and after the spermatogenesis to oogenesis switch (18, 19), generate oocytes or undergo programmed cell death. The proximal gonad contains a stack of oocytes, followed by sperms residing in the spermatheca. Both arms have a common uterus, where fertilized eggs are stored until hatching (20) (**Figure 1D**).

To visualize the germline, we dissected the gonads of Day 1 adult animals and stained them with DAPI. Confocal imaging showed that in *cdk-12* RNAi-fed *daf-2* worms, sperms were formed but oogenesis halted due to arrest of germ cells in the pachytene stage of meiosis. In *daf-16;daf-2*, the arrest was reversed and oocyte formation ensued (**Figure 1E**). Notably, upon *cdk-12* KD, the number of oocytes is reduced independent of DAF-16 (**Figure 1F**). The number of germ cells of *daf-2* in the pachytene stage of meiosis was drastically reduced upon *cdk-12* KD (**Figure 1G, H**); however, in *daf-16;daf-2* worms the reduction was largely abrogated (**Figure 1I, J**), leading to oocyte formation. The number of mitotic and transition zone nuclei remained unchanged in both cases (**Figure 1G-J**). We also found that the canonical IIS signalling pathway components are involved as *cdk-12* KD in *age-1(hx546)* (mutant in mammalian PI3K ortholog) (21) and *pdk-1(sa680)* (mutant in mammalian PDK ortholog) (22.)also arrested germline at the pachytene stage of meiosis **(Figure S1F, G)**.

DAF-16 has multiple isoforms with distinct and overlapping functions (10, 15, 23). We knocked down *cdk-12* in *daf-16;daf-2;daf-16a(+)* (DAF-16a rescued), *daf-16;daf-2;daf-16b(+)* (DAF-16b rescued) and *daf-16;daf-2;daf-16d/f(+)* (DAF-16d/f rescued) to find that the effect is mainly driven by DAF-16a (**Figure 1K, L**). Previously, DAF-16a isoform has been shown to play a major role in regulating lifespan, stress resistance and dauer formation (9, 15, 24, 25). Here, we show a predominant role of DAF-16a in preventing the pachytene exit of germ cells in *daf-2* when *cdk-12* is depleted.

### FOXO/DAF-16 and CDK-12 promotes a germline quality assurance program

Since the *daf-16;daf-2* worms produce oocytes when *cdk-12* is knocked down, in contrast to *daf-2*, we determined the quality of oocytes. As previously reported, we also found the oocytes produced on day 3 of adulthood by *daf-2* to be of superior quality in comparison to the wild-type worms (13). However, in *daf-16;daf-2,* the quality deteriorates significantly (Figure S2A-C) indicating the noted role of DAF-16 in the maintenance of better oocyte quality in *daf-2*. The quality of oocytes after *cdk-12* KD decreased in a DAF-16-independent manner (Figure 2A, B). It may also be noted that *cdk-12* KD decreases the number of hatched progenies in all strains; however, no brood is generated in *daf-2*. The brood size is partially rescued in *daf-16;daf-2* (Figure 2C).

**Figure 2.**
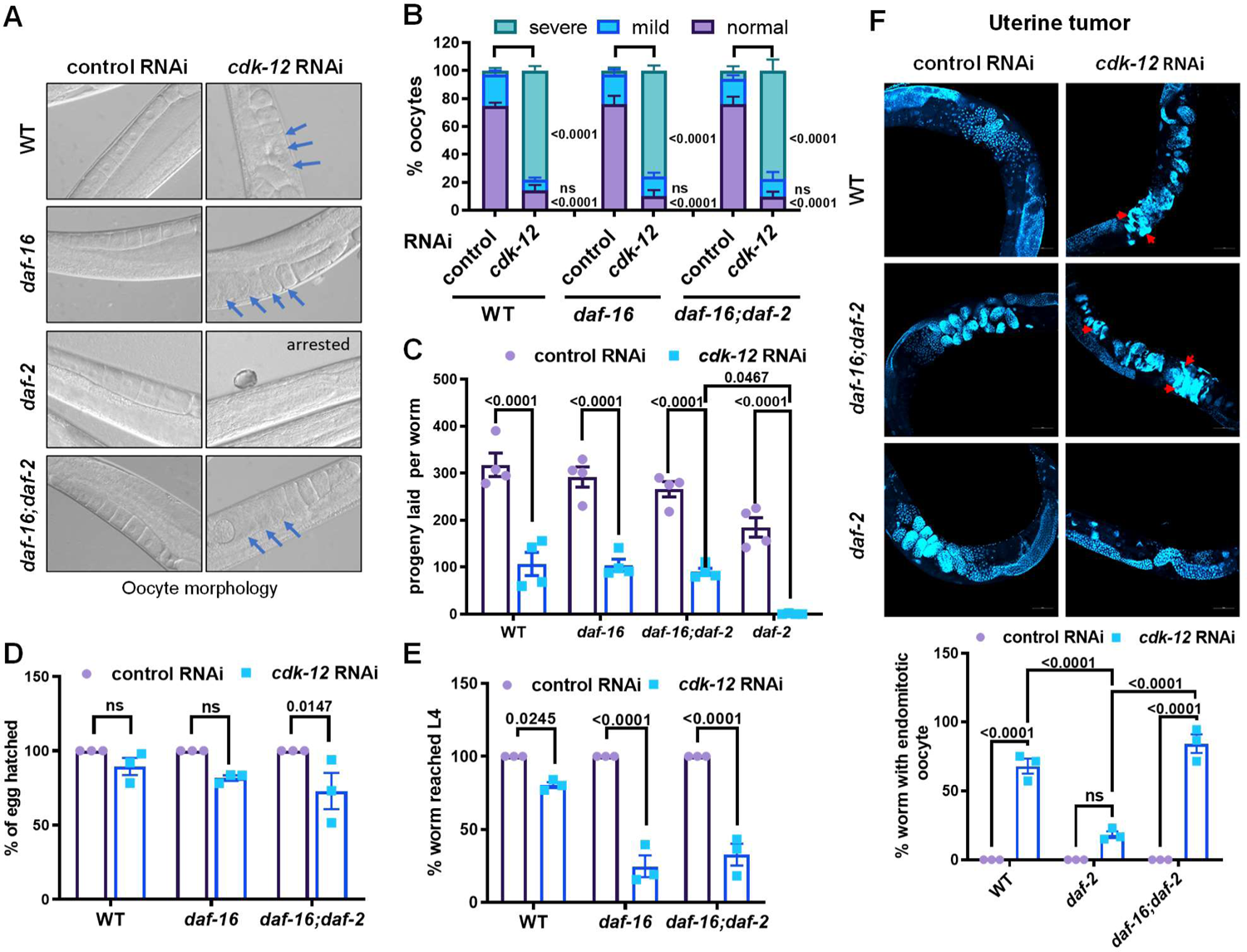
IIS pathway/*daf-16* and *cdk-12* regulate brood size, egg quality, oocyte quality and progeny health. (A) Representative DIC images of oocyte morphology when *cdk-12* was knocked down in WT, *daf-16(mgdf50), daf-2(e1370)* and *daf-16(mgdf50);daf-2(e1370)*. Blue arrows indicate morphologically disorganised oocyte. Image were captured at 400X magnification. (B) Quantification of oocyte quality on day 1 of adulthood in WT, *daf-16(mgdf50), daf-2(e1370)* and *daf-16(mgdf50);daf-2(e1370)* grown on control or *cdk-12* RNAi. The quality was categorized as normal, or with mild or severe defects according to images represented in Figure S2A. Average of four biological replicates (n≥25 for each replicate). One way ANOVA. (C) Lowered progeny count was observed in WT, *daf-16(mgdf50),* and *daf-16(mgdf50);daf-2(e1370)* on *cdk-12* RNAi, as compared to control RNAi. No progeny was observed in *daf-2(e1370)*. Average of four biological replicates (n≥14 for each replicate). Two-way ANOVA-Sidak multiple comparisons test. (D) Percentage of eggs that hatched in WT, *daf-16(mgdf50),* and *daf-16(mgdf50);daf-2(e1370)* upon *cdk-12* RNAi, as compared to control RNAi. Average of three biological replicates (n≥15 for each replicate). One way ANOVA. (E) The parental generation of different genetic background [WT, *daf-16(mgdf50)* or *daf-16(mgdf50);daf-2(e1370)*] was grown on *cdk-12* RNAi. The eggs were bleached and placed on control RNAi. Percentage of F1 that reached L4 or above after 72 hours is shown. Average of three biological replicates (n≥50 for each replicate). One way ANOVA. (F) Representative confocal images of worms stained with DAPI showing more endomitotic oocyte in WT, and *daf-16(mgdf50);daf-2(e1370)* as compared to *daf-2(e1370)* on *cdk-12* RNAi. The quantification of data is presented below. Average of three biological replicates (n≥10 for each replicate). Two-way ANOVA-Sidak multiple comparisons test. Image were captured at 240X magnification. Error bars are SEM. ns, non-significant. Experiments were performed at 20 °C. Source data is provided in Dataset S1.

Although most of the eggs that are laid by the *daf-16:daf-2* on *cdk-12* RNAi worms hatched (Figure 2D), they failed to reach the L4 stage (Figure 2E, S2D), indicating sub-optimal oocyte quality. Also, endomitotic oocytes (emo) that often develop due to defective fertilization (26), were more frequent in the proximal gonad of wild-type and *daf-16;daf-2* worms that were fed with *cdk-12* RNAi, compared to the *daf-2* worms (Figure 2F). Thus, we conclude that *cdk-12* plays an important role in maintaining oocyte quality and activated DAF-16, under conditions of lowered IIS, enforces a germline quality assurance program that prevents the production of inferior quality progeny.

### FOXO/DAF-16 and CDK-12 have shared transcriptional targets including DNA damage repair (DDR) genes

To understand the connection between the IIS pathway and CDK-12, we performed transcriptomics analysis at late L4 stage of WT, *daf-2* and *daf-16;daf-2* worm grown on control or *cdk-12* RNAi from L1 onwards. We found a large transcriptional response in *daf-2* when *cdk-12* is knocked down but not to that extent in WT (data not shown). Importantly, genes downregulated in *daf-2* on *cdk-12* RNAi are enriched for cell cycle, oogenesis, early embryonic development and hatching as well as DNA replication, repair processes (**Figure 3A**). When we compared the expression of the germline genes between *daf-2* and *daf-16;daf-2*, we found two distinct clusters, with one dependent on and the other independent of DAF-16 (**Figure 3B**). The fact that many important germline genes are downregulated in *daf-16;daf-2* supports our earlier observation that the quality of oocytes of the double mutant is poor. Out of the 4126 DAF-16-dependent genes upregulated in *daf-2*, 987 are also regulated by *cdk-12* **(Dataset S1)**, Similarly, out of the 1478 DAF-16-dependent genes downregulated in *daf-2*, 329 are *cdk-12* target, showing that DAF-16 and CDK-12 have shared transcriptional targets.

**Figure 3.**
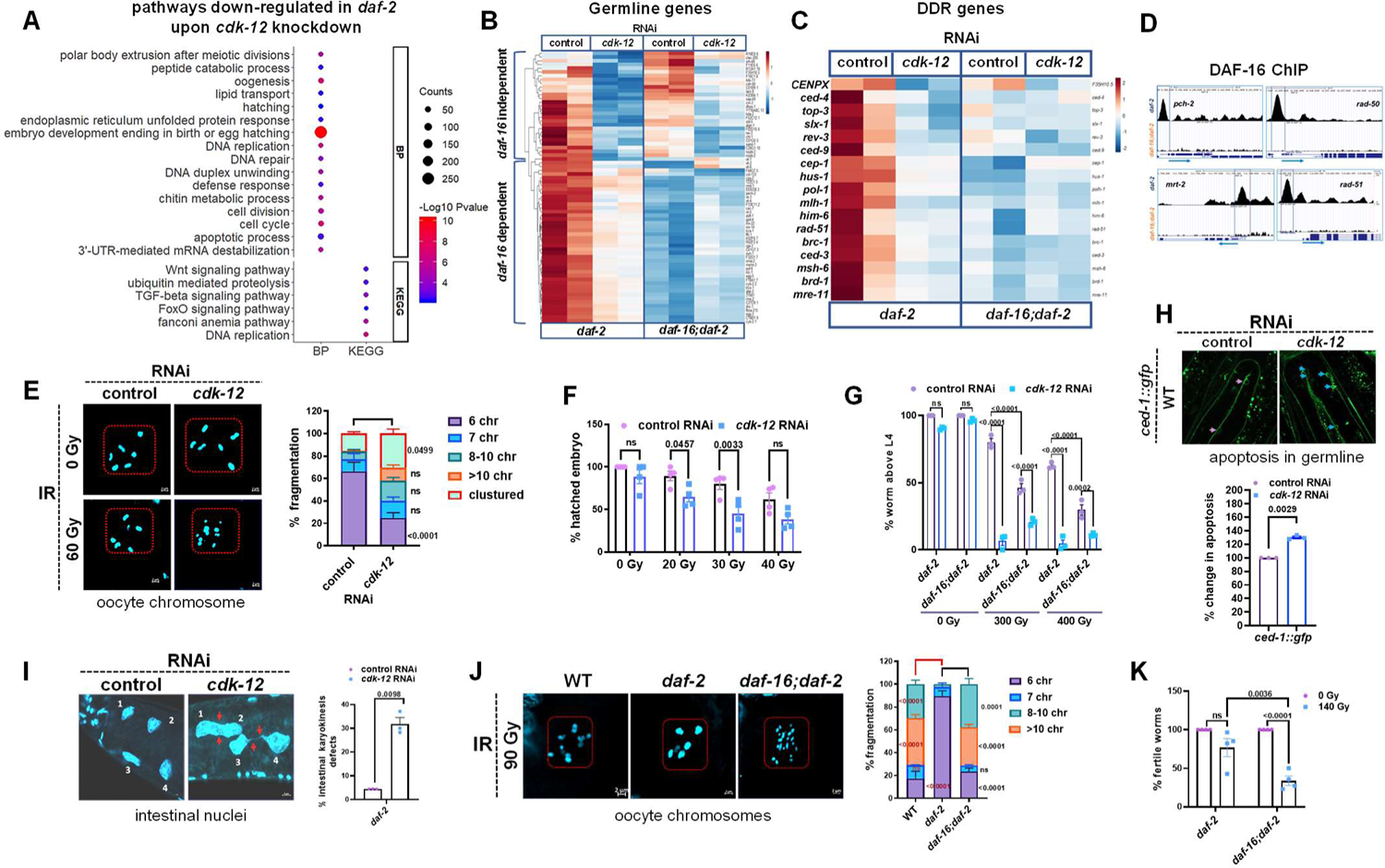
DAF-16 and CDK-12 regulate DDR gene expression for efficient DNA damage repair. (A) Gene Ontology (GO) Biological Processes (BP) term and KEGG pathway enrichment analysis of genes down regulated in *daf-2(e1370)* upon *cdk-12* KD using DAVID, as compared to control RNAi. (B) A heat map showing differential changes in the expression pattern of genes involved in germline development in *daf-2(e1370)* and *daf-16(mgdf50);daf-2(e1370)* upon control or *cdk-12* RNAi. (C) A heat map showing that DNA damage response (DDR) genes in *daf-2(e1370)* are down-regulated in a DAF-16-dependent manner. The DDR genes are also down-regulated in *daf-2(e1370)* upon cdk-12 RNAi, as compared to control RNAi. (D) UCSC browser view of FOXO/DAF-16 peaks on *pch-2, rad-50, mrt-2* and *rad-51* promoters as analysed by ChIP-seq analysis of *daf-2(e1370)* and *daf-16(mgdf50);daf-2(e1370)* strains. Blue boxes represent the promoter regions of *pch-2, rad-50, mrt-2* and *rad-51* having DAF-16 binding peaks in *daf-2(e1370)*. (E) Representative fluorescence images of DAPI-stained gonads showing oocytes with increased chromosome fragmentation upon γ-irradiation (60 Gy) in WT on *cdk-12* KD (left) and their quantification (right). Averages of four biological replicates (n≥59 oocyte for each replicate) are shown. One way ANOVA. (F) Decrease in the percentage of hatched embryo in WT grown on *cdk-12* RNAi upon γ-irradiation, as compared to control RNAi. Average of four biological replicates are shown (n≥20 for each replicate). One way ANOVA. (G) Worms of indicated strains were irradiated with different doses of γ-rays (0, 300, 400 Gy) at L1 larval stage and grown on control or *cdk-12* RNAi. After 96 hours, the percentage of worms that reached L4 or above was determined. Averages of 3 biological replicates (n≥100 for each replicate) are shown. Two-way ANOVA-Sidak multiple comparisons test. (H) Representative images showing apoptotic cell (arrow) in the gonadal arm of *ced-1::gfp* upon *cdk-12* KD and their quantification. Average of three biological replicates are shown (n≥17 for each replicate). Unpaired t test with Welch’s correction, Two-tailed. (I) Representative fluorescence images of DAPI-stained worms showing incomplete separation of intestinal cell nucleus upon *cdk-12* KD (left) and its quantification (right). Average of three biological replicates (n≥70 intestinal cell for each replicate). Unpaired t test with Welch’s correction, two-tailed. (J) Representative DAPI-stained fluorescence images of oocytes showing chromosome fragmentation in WT, *daf-2(e1370)* and *daf-16(mgdf50);daf-2(e1370)* upon treatment with γ-irradiation (90 Gy) (left) and their quantification (right). Average of two biological replicates (n≥27 for each replicate) is shown. One way ANOVA. (K) The *daf-2(e1370)* and *daf-16(mgdf50);daf-2(e1370)* worms were exposed to γ-irradiation (140 Gy) at L1 larval stage. Quantification showing percentage fertile worms. Arrows indicate sterile worms. Average of three biological replicates (n≥20 for each replicate) are shown. Two-way ANOVA-Sidak multiple comparisons test. Error bars are SEM. ns, non-significant. Experiments were performed at 20 °C. Source data is provided in Dataset S1.

In mammalian cells, CDK12 specifically regulates genes involved in DNA damage response (17, 27). We also found the DDR gene expression in *daf-2* to be considerably down-regulated upon *cdk-12* KD. In addition, these genes were also dependent on DAF-16 (**Figure 3C**).

We validated this by quantitative real-time PCR (RT-PCR) **(Figure S3A)**. We also found many of these genes to be down-regulated in WT on *cdk-12* RNAi **(Figure S3B)**. The downregulation in *daf-2* was not due to differences in the sizes of the gonads upon *cdk-12* KD **(Figure S3C)**. Importantly, ChIP-seq data analysis in *daf-2* and *daf-16;daf-2* showed that many DNA damage checkpoint genes like *mrt-2*, *rad-51*, *rad-50* and *pch-2* and DNA damage repair and cell cycle genes have DAF-16 binding peaks in their promoter-proximal regions (**Figure 3D**, S3D). suggesting that they may be direct targets of DAF-16. Together, these data show that genes involved in sensing and repairing DNA damage are common transcriptional targets of DAF-16 and CDK-12.

### CDK-12 is required for efficient DNA damage repair

Mammalian CDK12 is known to regulate DDR genes and promote homologous recombination (HR)-mediated DNA repair (17, 28). Above, we also found CDK-12 to transcriptionally regulate the DDR genes in *C. elegans*. To determine whether CDK-12 KD leads to germline DNA damage, we utilized a chromosome fragmentation assay. In unirradiated worms, six highly condensed bivalent bodies can be seen in the oocyte; however, unrepaired DNA strand breaks in irradiated worms lead to chromosome fragmentation/fusions (29). We observed increased chromosome fragmentation and fusions in IR-treated wild-type worms upon *cdk-12* KD that suggests increased DNA damage (**Figure 3E**). Next, we exposed the L4 or YA worms to different concentrations of DNA damaging agent camptothecin (CPT) **(Figure S3E)** or varying doses of Ionizing Radiation (IR) (**Figure 3F**) and found that *cdk-12* KD resulted in a lesser number of hatched eggs, highlighting their higher sensitivity, possibly due to compromised DNA damage repair. We also observed increased developmental arrest on IR treatment at L1 stage when *cdk-12* is KD, in a DAF-16-independent manner (**Figure 3G**).

In agreement to the fact that *cdk-12* KD may lead to endogenous DNA damage, we observed higher apoptotic bodies per gonadal arm in *cdk-12* KD wild-type and *daf-2* worms (**Figure 3H**). DNA damage in worm germline has been shown to evoke the innate immune response which in turn confers systemic resistance and enhances somatic stress endurance (30). In our transcriptomic data, we find that KD of *cdk-12* up-regulates innate immune response genes independent of DAF-16 activation **(Figure S3F, G)**. Further, *cdk-12* depletion conferred increased heat stress resistance **(Figure S3H, I)** and *hsp-4::gfp* (Endoplasmic Reticulum Chaperon BiP ortholog) expression **(Figure S3J)**, as has been reported for DNA damage (30, 31). Thus, RNAi depletion of *cdk-12* may cause DNA damage in cells that may be sensed by DAF-16 in the *daf-2* mutant.

Further, we wanted to visualize the role of CDK-12 in somatic DNA damage. For this, we analysed the DAPI-stained adult intestinal cells. A total of 20 intestinal cells are present at hatching, a subset of which (8–12) divide, but do not undergo cytokinesis, thereby generating 28-32 binucleate intestinal cells by the end of the L1 stage (32). Like mutations in some DDR genes, *atm-1* and *dog-1* (33), we also found elongated cells with chromosomal bridges upon *cdk-12* KD (**Figure 3I**), much similar to L4 worms exposed to IR **(Figure S3K),** indicating the occurrence of DNA damage in the somatic cells (29, 33).

Together, CDK-12 plays a pivotal role in the repair of damaged DNA, both in the *C. elegans* germline and somatic tissues to maintain genomic integrity. Therefore, knocking down *cdk-12* may lead to genomic instability that is sensed by activated DAF-16 in the *daf-2* mutant, leading to the germline arrest at pachytene stage of meiosis. The DNA damage on *cdk-12* KD also accelerates aging independent of DAF-16.

### FOXO/DAF-16 confers increased DNA damage repair efficiency

To test the functional role of DAF-16 in DDR and its heightened engagement in *daf-2* to protect against DNA damage, we again utilized the chromosome fragmentation assay. Worms were treated with IR at L4 to induce DNA double-strand breaks, stained with DAPI after 48 hours post-radiation and imaged. We found *daf-2* worms to be highly resistant to IR, such that at 90 Gy most of the wild-type chromosomes were fragmented, but *daf-2* worms retained intact chromosomes. This IR resistance was conferred by DAF-16, as in the *daf-16;daf-2* worms, the chromosomes were fragmented to a similar extent as in wild-type with IR treatment (**Figure 3J**).

A high dose of gamma radiation during early larval stages in *C. elegans* can result in sterility and developmental arrest if the damage is not repaired (34). Upon treatment of *daf-2* and *daf-16;daf-2* worms with 140 Gy IR dose at the L1 stage, we found that *daf-16;daf-2* worms become sterile (**Figure 3K**). However, remarkably, *daf-2* worms were mostly fertile. Similarly, resistance to somatic developmental arrest on IR treatment was observed in *daf-2*, in a *daf-16-*dependent manner (**Figure 3G**). Together, our findings support a role of DAF-16 in regulating DNA damage repair during lowered IIS, thereby promoting resistance to DNA damage, supporting growth and reproduction.

### Regulation of pachytene arrest in *daf-2* upon DDR perturbation

IIS pathway couples nutrient sensing to meiosis progression and oocyte development to enable reproduction only when conditions are favourable for survival (2). The well conserved LET-60 (RAS)-MEK-2 (ERK kinase)-MPK-1 (ERK1/2) pathway has several roles in the germline development and its maturation (35). The RAS-ERK pathway works downstream of the IIS receptor *daf-2*. In response to nutrient availability, IIS activates MPK-1 (ERK) to promote meiotic progression. Thus, in the absence of nutrients or low food conditions, MPK-1 inhibition results in stalling of meiosis. In the *daf-2* germline stained with pMPK-1 antibody, the level of ERK activation is significantly lower than WT (36) (**Figure 4A, B**). This potentially explains why *daf-2* have reduced brood size and oocyte numbers (**Figure 2C**, S4A). This level is rescued to WT levels in *daf-16;daf-2* worms, showing that DAF-16 may negatively regulate pMPK-1 levels (**Figure 4A, B**). When *cdk-12* is knocked down in WT, the levels of pMPK-1 is significantly reduced. However, the reduction is much more dramatic in *daf-2*, possibly below a threshold level (**Figure 4A, B**). This may explain the complete arrest of the germline at pachytene stage (**Figure 1E**). Importantly, in the *daf-16;daf-2*, the levels are restored (**Figure 4A, B**), in line with the release of pachytene arrest in the double mutant (**Figure 1E**).

**Figure 4.**
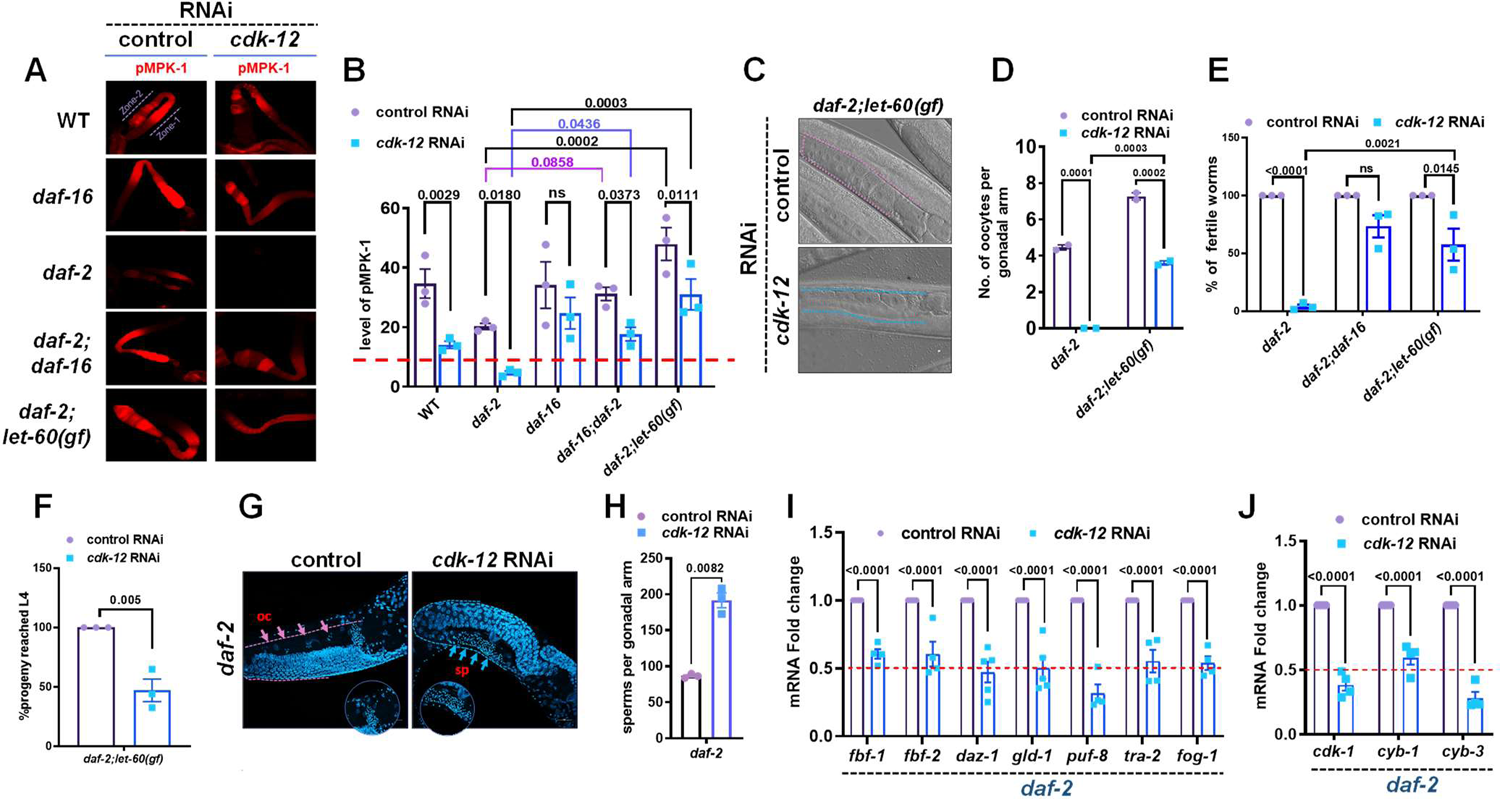
CDK-12 KD disturbs the balance of ERK-MPK and IIS signaling that regulates germline development. (A-B) Representative images of dissected gonads of WT, *daf-2(e1370), daf-16(mgdf50);daf-2(e1370)*, *daf-16(mgdf50)* and *daf-2(e1370);let-60(ga89)*, probed with anti-dpERK (red) (A) and its quantification (B) upon control or *cdk-12* RNAi. Average of three biological replicates (n≥10 for each replicate). Two-way ANOVA-Uncorrected Fisher’s LSD multiple comparisons test. Zone-1 and zone-2 are the proximal and distal parts of the gonad, respectively. Red line in (B) is a presumptive threshold of pMPK-1 below which germline arrests. (C) Representative DIC images of worms showing oocytes of *daf-2(e1370);let-60(ga89)* upon *cdk-12* KD. (D) Quantification of oocyte number per gonadal arm of *daf-2(e1370)* and *daf-2(e1370);let-60(ga89)* upon *cdk-12* KD. Averages of two biological replicates (n≥20 for each replicate). Two-way ANOVA-Sidak multiple comparisons test. (E) Percentage of fertile worms in *daf-2(e1370)*, *daf-16(mgdf50);daf-2(e1370)* and *daf-2(e1370);let-60(ga89)* on control or *cdk-12* RNAi. Average of three biological replicates (n≥25 for each replicateTwo-way ANOVA-Sidak multiple comparisons test. The concentration of IPTG used in this experiment is 0.4mM. (F) The *daf-2(e1370);let-60(ga89)* were grown on control or *cdk-12* RNAi. The worms were bleached and their eggs grown on control RNAi. Percentage of hatched progeny that reached L4 larval stage is plotted. Average of three biological replicates (n≥40 for each replicate). Unpaired t test with Welch’s correction, Two-tailed. (G, H) DAPI stained worms and quantification of the sperm count in *daf-2(e1370)* on control and *cdk-12* RNAi. Average of three biological replicates (n≈20 for each replicate). Unpaired t test with Welch’s correction, Two-tailed. (I) Quantitative RT-PCR analysis of sperm-to-oocyte switch genes in *daf-2(e1370)* on control or *cdk-12* RNAi. Expression levels were normalized to *actin*. Average of four biological replicates are shown. One way ANOVA (J) Quantitative RT-PCR analysis of cell cycle regulator *cdk-1* and its binding partner *cyb-1* and *cyb-3* (mammalian Cyclin B orthologs) in *daf-2(e1370)* upon *cdk-12* KD, compared to control RNAi. Expression levels were normalized to *actin*. Average of four biological replicates are shown. One way ANOVA. Error bars are SEM. ns, non-significant. Experiments were performed at 20 °C. Source data is provided in Dataset S1.

It appears that downstream of *daf-2*, the ERK signalling and the canonical PI3K signalling co-ordinately regulate germline pachytene arrest. When *daf-2* is mutated, the pMPK-1 levels are lowered because of less signalling through the RAS pathway as well as due to the negative regulation of activated DAF-16 through the PI3K pathway. We have shown above that knocking out *daf-16* rescues the lower pMPK-1 in *daf-2* (**Figure 4A, B**). So, we asked whether activating the ERK signalling can bypass the pachytene arrest in *daf-2* on *cdk-12* KD. We used an activated *ras* allele with constitutively high pMPK-1 phosphorylation (37). In the *daf-2;let-60(gf)*, the pMPK-1 levels were upregulated (**Figure 4A, B**) and pachytene arrest was partially reversed (**Figure 4C-E**). Although many eggs hatched to release L1 worms **(Figure S4B),** only about half of them were able to reach adulthood (**Figure 4F**), possibly pointing at their poor quality. Overall, we conclude that the ERK and the PI3 kinase pathways co-ordinately regulate meiosis arrest on sensing somatic DDR perturbations in *daf-2*.

### Defective sperm to oogenesis switch and transcriptional downregulation of key cell cycle genes in *daf-2* on DDR perturbation

We have shown above that the sterility of *daf-2* on *cdk-12* RNAi may be due to inactive RAS-ERK signaling. RAS-ERK activation is critical for sperm-oocyte fate switch by regulating the timing the event in *C. elegans* hermaphrodite(38). We observed a two-folds increase in the number of sperms but no oocyte in *daf-2* upon *cdk-12* KD (**Figure 4G, H**). So, we tested the mRNA levels of key sperm-oocyte switch genes and found their levels to be significantly reduced in *daf-2* (**Figure 4I**, but not in *daf-16;daf-2* **(Figure S4C)**. This decrease in expression of genes is due to the *cdk-12* KD *per se*, and not because of a reduction in germline size as at late-L4 (when RNA was collected), the germline size is comparable between control RNAi and *cdk-12* RNAi fed worms **(Figure S3C).**

Next, we asked if the sperm to oogenesis switch defect was accompanied by an underlying defect in other critical players of meiotic progression, namely, *cdk-1*, *cyb-1* and *cyb-3* (39). To assess this, we determined the mRNA levels of these genes and found levels of all three to be significantly down-regulated in *daf-2* worms with *cdk-12* KD (**Figure 4J**), whereas the gene levels were largely unchanged in *daf-16;daf-2 cdk-12* RNAi worms **(Figure S4D)**. Additionally, knocking down these genes individually led to sterility in *daf-2* worms **(Figure S4E, F)**, phenocopying the sterility upon *cdk-12* KD.

We further checked if a similar defect in sperm to oogenesis switch and downregulation of key cell cycle genes underlies the sterility upon DNA damage on IR exposure. We treated *daf-2* worms with 160 Gy IR at L1 and DAPI stained Day 1 adults. Surprisingly, we found that the sperm count increased around two-fold with a concomitant reduction in sperm to oocyte switch genes and *cdk-1*, *cyb-1*, and *cyb-3* RNA levels **(Figure S4G-I)**. Therefore, using CDK-12 knockdown and IR exposure to phenocopy DNA damage, we show that germline arrest on DDR perturbation in *daf-2* is brought about by defective sperm to oogenesis switching and reduction in the transcription of genes essential for meiotic progression. This, along with reduction of ERK/MPK-1 signaling, may be strategies employed by the *daf-2* hermaphrodite worms to prevent the production of poor-quality progeny when the DNA damage is beyond repair.

### Uterine tissue-specific DDR perturbation arrests germline in *daf-2*

Since *cdk-12* KD leads to impaired DDR and resulting DNA damage, we asked whether tissue-restricted DNA repair perturbations will lead to germline arrest in *daf-2*. We first used a germline-specific RNAi system to test if tissue autonomous depletion was sufficient for the arrest in *daf-2*. We used a *rde-1*(*-*) transgenic strain where *sun-1* promoter drives the expression of *rde-1* only in the germline of *daf-2* (germline-specific RNAi) (40). We validated the strain by knocking down a germline-specific gene *glp-1* (41) which led to sterility **(Figure S5A)**, showing a functional germline RNAi machinery. A systemic KD of a soma-specific GATA transcription factor, *elt-2* (42) leads to developmental arrest in wild-type; however, the *daf-2* germline-specific RNAi worms were resistant to *elt-2* KD **(Figure S5A)**, showing the lack of RNAi in the somatic tissues. Surprisingly, we found KD of *cdk-12* only in germline does not lead to sterility (**Figure 5A, B**), indicating that a soma-specific DDR malfunction may cause the germline arrest. However, depletion of *cdk-12* in the germline alone results in progenies that are developmentally arrested and sterile (**Figure 5C**), showing that its function is required in the germline to maintain progeny quality. Importantly, it appears that activated DAF-16 only promotes germline arrest if the damage signal emanates from somatic tissues.

**Figure 5.**
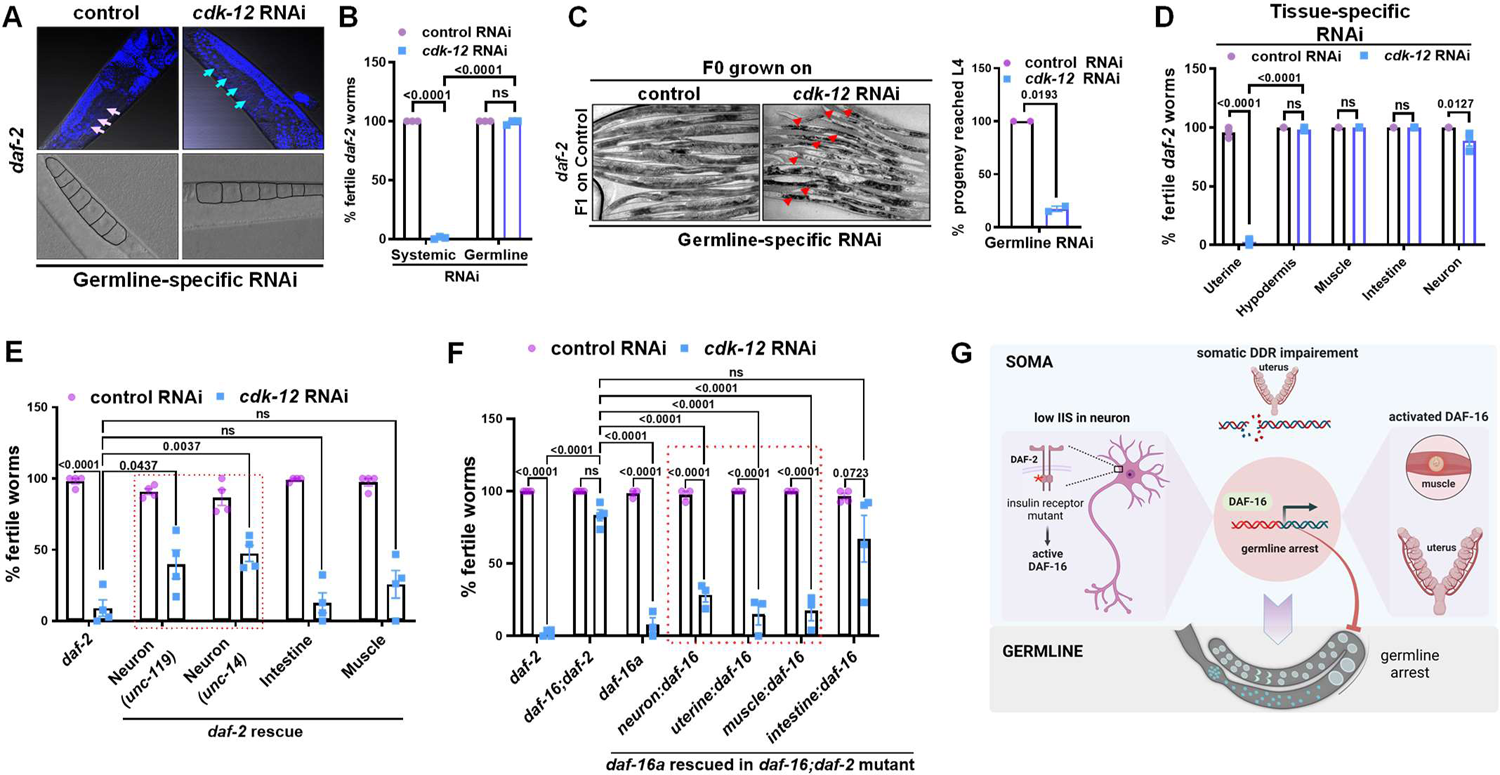
Cell non-autonomous signals from soma determines germline fate. (A) The *cdk-12* knock down by RNAi specifically in the germline of *daf-2(e1370);mkcSi13 II; rde-1(mkc36) V [mkcSi13 [sun-1p::rde-1::sun-1 3’UTR + unc-119(+)] II]* (germline-specific RNAi) strain produced no arrest. Arrows showing oocyte nuclei. Upper panels are 400x DAPI images and lower panels are bright field. (B) Percentage of fertile worms in *daf-2(e1370)* and *daf-2(e1370);mkcSi13 II; rde-1(mkc36) V [mkcSi13 [sun-1p::rde-1::sun-1 3’UTR + unc-119(+)] II]* (germline-specific RNAi) on control or *cdk-12* RNAi. Average of three biological replicates (n≥30 for each replicate). Two-way ANOVA-Sidak multiple comparisons test. (C) *Cdk-12* was knocked down specifically in the germline of *daf-2(e1370)* using germline-specific RNAi strain *(daf-2(e1370);mkcSi13 II; rde-1(mkc36) V [mkcSi13 [sun-1p::rde-1::sun-1 3’UTR + unc-119(+)] II])*. The eggs produced by these worms or the ones grown on control RNAi were transferred to fresh control RNAi plates. Representative brightfield images of these F1 progeny shown along with quantification. Average of two biological replicates (n≥30 for each replicate). Unpaired t test with Welch’s correction, Two-tailed. (D) Percentage of fertile worms when *cdk-12* is knocked down in different tissues of *daf-2(e1370)*. Only uterine-specific knockdown [using *daf-2(e1370);rrf-3(pk1426) II; unc-119(ed4) III; rde-1(ne219) V; qyIs102*] of *cdk-12* results in sterility. Average of three biological replicates (n≥19 for each replicate). Error bars are SEM. One way ANOVA. (E) Percentage of fertile worms on *cdk-12* KD in strains where the *daf-2* gene is rescued either in the neurons, intestine or muscles of the *daf-2(e1370)* mutant. Average of four biological replicates (n≥30 for each replicate). Two-way ANOVA-Sidak multiple comparisons test. (F) Percentage of fertile worms on *cdk-12* KD in strains where the *daf-16* gene is rescued either in the neurons, intestine, muscles or uterine tissues of the *daf-2(mu86)* mutant. Average of four biological replicates (n≥15 for each replicate). Two-way ANOVA-Sidak multiple comparisons test. (G) A tentative model showing inter-tissue crosstalk of low IIS in the neuron and activated DAF-16/FOXO in the neuron or muscle or uterine tissue (somatic gonad) that is required to mediate the germline arrest in response to somatic DNA damage or DDR perturbation by *cdk-12* depletion. Error bars are SEM. ns, non-significant. Experiments were performed at 20 °C. Source data is provided in Dataset S1.

**Figure 6.**
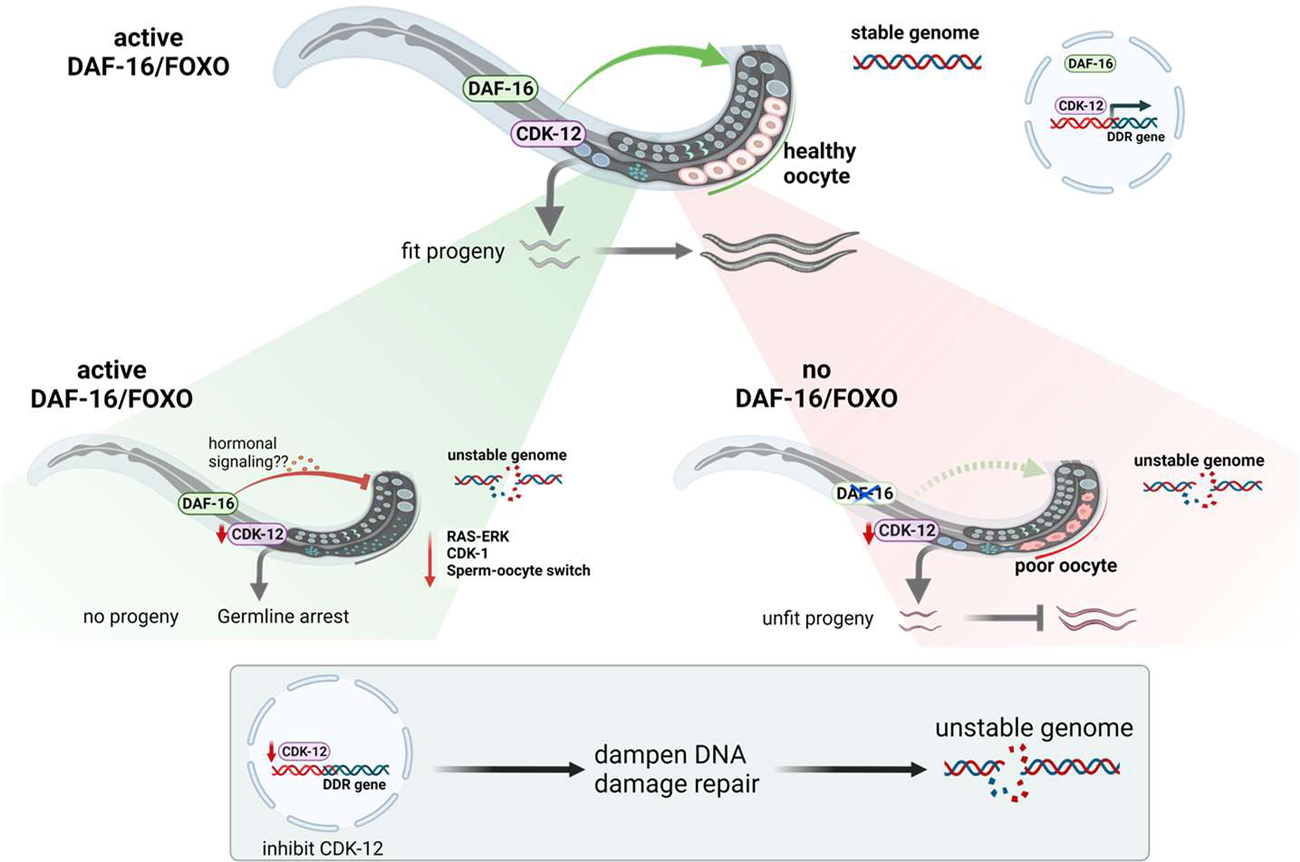
A model showing how activated DAF-16/FOXO may act as a quality control checkpoint from the somatic tissue, regulating reproductive decision, gamete quality and progeny health, likely via soma-germline hormonal signaling. Somatic DAF-16/FOXO may sense the DDR inactivation and send signals to germline to halt reproduction (by germline arrest) in order to protect the genome integrity of the germ cells/oocytes and maintain progeny fitness. In the absence of active DAF-16/FOXO, organisms fail to arrest reproduction and produces compromised oocyte and unhealthy progeny. DDR perturbation only in the somatic tissue (uterus) is sufficient to arrest the germline. DAF-16/FOXO arrests the germ cell development by inactivating the RAS-ERK signaling which is essential for germline proliferation and its maturation.

Next, we specifically knocked down *cdk-12* in different somatic tissues (43–46). We found that knocking down *cdk-12* only in the uterine tissues was sufficient to arrest the germline in the *daf-2* worms at the pachytene stage of meiosis (**Figure 5D**, S5B); no arrest was seen when the gene was knocked down in hypodermis, muscle, intestine or neurons (**Figure 5D**) and they produced healthy fertile progeny **(Figure S5C)**. This implies that KD of *cdk-12* in *daf-2* germ cells may lead to DNA damage resulting in poor progeny production. However, knocking it down in the somatic uterine tissue may activate DAF-16-dependent quality checkpoints that lead to cell-nonautonomous germline arrest.

Next, to determine the tissues where the IIS receptor functions, we used transgenic lines where the wild-type copy of *daf-2* is rescued only in the neurons (using either *unc-119* or *unc-14* promoters), muscles or intestine of the *daf-2* mutants (47) and then knocked down *cdk-12* using RNAi. We found that neuron-specific rescue of the *daf-2* gene led to a significant rescue of fertility, while little effect was seen in the case of muscle and intestine-specific rescue (**Figure 5E**). We also determined where DAF-16 is required to sense and mediate the germline arrest in the *daf-2* mutant upon *cdk-12* KD. We found maximum arrest when *daf-16* is rescued in the muscle, neuron or uterine tissues of the *daf-16;daf-2* mutant worms (**Figure 5F**), but not in the intestine. Together, these observations support a model where low neuronal IIS sensitizes uterine tissues to perturbations in DDR, leading to the arrest of germline at the pachytene stage of meiosis. The DAF-16a isoform works in the somatic uterine tissues, apart from muscle and neurons, to implement the arrest (**Figure 5G**).

## Discussion

In this study, we have shown that activated FOXO/DAF-16 senses the intrinsic somatic DNA damage and functions cell non-autonomously to regulate reproductive decision in order to safeguard the germline genomic integrity and progeny fitness.

CDK-12 is a well-studied protein that is involved in DDR and genome integrity in mammalian cells. We also show that in *C. elegans*, *cdk-12* regulates the expression of DDR genes and maintains genome integrity (48, 49). The depletion of *cdk-12* makes the worms susceptible to DNA damaging agents and induce spontaneous DNA damage both in the soma and the germline, implying a suboptimal repair pathway. We show *cdk-12* ablation reduces gamete quality, leading to increased infertility and decreased progeny fitness. It also led to retarded growth, and premature aging which are hallmarks of genomic instability. Thus, CDK-12 is an evolutionary well-conserved custodian of the genome that helps maintain DNA integrity, which we have used in our study as a genetic tool to analyse the effects of tissue-restrictive DDR perturbation and DNA damage.

To maintain genomic integrity, organisms have evolved an efficient DDR pathways, that senses and repairs DNA damage (50). Defects in DDR is associated with reduced fitness, infertility and offspring with inherited diseases (51, 52). We identified that active FOXO/DAF-16 maintains genomic integrity by upregulating DNA repair genes, which could explain the longer lifespan and better oocyte quality of the low IIS mutants (53). Apart from maintaining genomic stability, our study shows that activated FOXO/DAF-16 can sense DDR perturbation or DNA damage, stop reproduction by arresting the germline, and protect the genomic integrity of germ cell. In the absence of DAF-16, worms fail to arrest germline development and produce oocytes of poor quality that hatch into unhealthy progenies. Thus, activated FOXO/DAF-16 critically regulates reproductive decision by sensing the intrinsic threat of genomic instability. Previous studies has shown that DAF-16 acts as a nutrient sensor and mediates developmental arrest on starvation, as a protective mechanism (7). Together, these data suggests that FOXO/DAF-16 acts as a master regulator of diverse cellular processes in maintaining genomic integrity, tissue homeostasis and reproduction.

We found that upon DDR perturbation and ensuing DNA damage, FOXO/DAF-16 enforces germline arrest by inactivating RAS-ERK signalling which is essential for germline proliferation and quality. We also observed reduced expression of cyclin-dependent kinase-1 gene (*cdk-1*) and its binding partner cyclin, *cyb-1,* and *cyb-3* genes, which may be due to dampening of the RAS-ERK signaling. In many cancers, RAS-ERK negatively regulates FOXO activity and promotes rapid proliferation(54). Similarly, we observed that constitutively activated RAS-ERK in the low IIS mutant (where FOXO/DAF-16 is activated) over-rides the germline arrest upon DDR perturbation, leading to the production of unhealthy progenies. Therefore, RAS-ERK and FOXO/DAF-16 regulate each other’s activity and a fine balance is important for various biological process, including reproductive development.

Cell non-autonomous inter-tissue crosstalk helps an organism to perceive and respond to changing environment. Multiple studies in *C. elegans* have revealed cell non-autonomous crosstalk in stress response and longevity (55). DAF-2 in the neuron and DAF-16 in the intestine is known to regulate longevity cell non-autonomously (14, 15). Muscle or intestinal FOXO/DAF-16 activity promotes long reproductive span or better oocyte quality of the *daf-2* mutant (13). DAF-16 has also been shown to function in the uterine tissue to prevent decline in the germline progenitor cells with age (11). However, it is not clear how DAF-16 cell non-autonomously regulates germline health. We show that perturbation of the DDR pathway only in the somatic uterine tissue of low IIS worm is sufficient to cause cell cycle arrest in the germline; perturbation in the germline itself does not lead to arrest but produces unhealthy progenies. This suggests that the somatic tissue, not the germline, senses stress signal of genome instability and shunt their energy and resources towards somatic maintenance rather than reproductive commitment. This is supported by the observations of heightened stress response pathways and retarded germline growth upon DDR perturbation.

Finally, we find that lowering of IIS is required in the neurons to activate FOXO/DAF-16 cell-autonomously in the neurons as well as non-autonomous in the muscle and uterine tissues to mediate cell cycle arrest. This soma to germline communication is most likely mediated by the DAF-12/dafachronic acid steroid signalling. It has been previously shown that DAF-12/dafachronic acid steroid signalling is required for germline to soma signalling for DAF-16-dependent longevity in germline mutants (56, 57) Previous studies have also shown that neurons can sense cell intrinsic unfolded protein stress and mount a protective response in distal tissues (58). Thus, from an evolutionary perspective, such a complex network of non-autonomous inter-tissue crosstalk likely helps an organism sense intrinsic or extrinsic stresses more efficiently and accurately. This may ensure the optimal survival as well as fitness across generations.

## Materials and Methods

Complete materials and methods are provided in SI Appendix file.

### DAPI staining

Worms were grown on control or *cdk-12* RNAi, L1 stage onwards. Day 1 adults were collected in 1X M9 buffer in a 1.5 ml Eppendorf tube and worms were allowed to settle down. Using a glass Pasteur pipette, the 1X M9 was discarded, leaving behind a ∼ 100 µl worm suspension. Then, 1 ml chilled 100 % methanol was added to the worm pellet and incubated for 30 minutes at −20°C. The pellet was placed on a glass slide and Fluroshield with DAPI (Invitrogen, Carlsbad, USA) was added. For staining dissected gonads, worms were placed onto a glass slide and the gonads were obtained by cutting the pharynx or tail end of the worm using a sharp 25G needle. After collecting gonads for 10 minutes, 500 µl chilled 100% methanol was added onto the slide and allowed to dry. Fluroshield with DAPI (Invitrogen, Carlsbad, USA) was added. The slides were imaged using a confocal microscope (Carl Zeiss, Oberkochen, Germany).

### Reproductive span, brood size, and egg hatching

Worms were grown on control or *cdk-12* RNAi from L1 onwards and upon reaching the young adult stage, five worms were picked onto fresh RNAi plates, in triplicates, and allowed to lay eggs for 24 hours. The worms were then transferred to fresh plates every day until worms ceased to lay eggs, and the eggs laid on the previous day’s plate were counted. These plates were again counted after 48 hours to document the number of hatched worms and the un-hatched eggs were considered dead eggs. The pool of hatched and dead eggs is defined as brood size. Egg quality was determined by calculating the percentage of hatched progeny in different conditions.

### Assay to quantify developmental retardation of progeny

Synchronized L1 worms were grown on different RNAi and allowed to lay eggs. The eggs and L1s were transferred to control RNAi plates. After 72 hours on control RNAi, the plates were scored for progeny that reached the L4 stage or beyond.

### Analysis of Oocyte Morphology

Worms were grown from L1 onwards on control or *cdk-12* RNAi. Differential interference contrast (DIC) images of the oocytes were captured on day 1 and day 3 of adulthood. Oocyte images were categorized into three groups based on their morphology (cavities, shape, size, and organization). Based on the severity of the phenotype, oocytes were categorized as normal (no small oocytes, no cavities, and no disorganized oocytes), mild (a few cavities in gonad, or slightly disorganized oocyte or small), or severe (many cavities in the gonad, or disorganized or misshapen).

### Quantification of fertile worms

Worms were bleached and their eggs were allowed to hatch in 1X M9 buffer for 17 hours to obtain L1 synchronized worms. Approximately 100 L1 worms were placed onto different RNAi plates, in triplicates. On day 1 adult stage, bright-field images were captured (Carl Zeiss, Oberkochen, Germany). Worms with more than five eggs in the uterus were considered fertile.

### Oocyte number

Approximately 100 L1 worms were placed onto control or *cdk-12* RNAi plates, in triplicates. On day 1 adult stage, DIC Image of oocyte was captured (Carl Zeiss, Oberkochen, Germany) and the oocyte number per gonadal arm was counted.

### Chromosomal fragmentation assay

Approximately 100 L1 worms were placed onto control or *cdk-12* RNAi plates, in triplicates. The worms were irradiated with ionizing radiation (IR) of different doses at the L4 stage. After 48 hours, the worms were stained with DAPI and the oocyte chromosomes were imaged in Z-stack using an LSM980 confocal microscope (Carl Zeiss, Oberkochen, Germany). For scoring chromosome fragmentation, images were converted into maximum intensity projection (MIP) and scored.

### Scoring of Endomitotic oocyte

Approximately 50 L1 worms were placed onto control or *cdk-12* RNAi plates, in triplicates. Day 1 adult worms were stained with DAPI and imaged in Z-stack using an LSM980 confocal microscope (Carl Zeiss, Oberkochen, Germany). For scoring, the images were converted into maximum intensity projection (MIP) and scored.

### Intestinal cell nucleus morphology

Approximately 50-80 L1 worms were placed onto control or *cdk-12* RNAi plates, in triplicates. Day 1 adult worms were stained with DAPI and the intestinal nucleus was imaged using an LSM980 confocal microscope (Carl Zeiss, Oberkochen, Germany).

### Ionizing radiation (IR) treatment of Larval stage 1 worms

For sterility assay, approximately 100 L1 worms were placed onto control RNAi plates and treated with different doses of IR. Day 1 adult worms were imaged under a bright-field microscope (Carl Zeiss, Oberkochen, Germany). Worms with more than five eggs in the uterus were considered to be fertile.

Similarly for developmental assay, approximately 100-130 L1 worms were treated with different doses of IR on control RNAi. Then, IR-treated L1 were transferred to different RNAi plates and scored for worms that reached L-4 or above post 100 hours.

### DNA damage sensitivity assay

#### IR

Worms were grown on control or *cdk-12* RNAi. At the young adult stage, worms were exposed to IR doses ranging between 0 to 40 Gy. The IR-treated worms were allowed to recover for 3-4 hours, following which 5 worms were transferred to respective RNAi plates, in duplicates, and then incubated for 18–20 h at 20 °C. The adults were sacrificed and the number of eggs laid on the plates was counted. About 48 hours later, the number of hatched progenies was also counted.

#### Camptothecin

The working stock of CPT (2 µM) was made in 10 X concentrated bacterial feed suspended in 1x M9 buffer. Worms were added to wells containing CPT in liquid bacterial feed. The plates were wrapped with foil and incubated at 20 °C for 18–20 h, with gentle shaking. The worms were then transferred to Eppendorf tubes and washed twice with 10% Triton X-100 (in 1x M9 buffer), followed by two washes with 1x M9 buffer. The worms were then placed on RNAi plates to recover for 3-4 hours, followed by tight egg-laying for 3-4 hours on fresh, respective RNAi plates. The adults were sacrificed and the number of eggs laid was counted. About 48 hours later, the number of hatched progenies was also counted.

## Acknowledgments

We thank all members of Molecular Aging for their support. This project was partly funded by National Bioscience Award for Career Development (BT/HRD/NBA/38/04/2016), SERB-STAR award (STR/2019/000064), DBT grants (BT/PR27603/GET/119/267/2018; BT/PR16823/NER/95/304/2015), ICMR grant (54/3/CFP/GER/2011-NCD-II) and core funding from the National Institute of Immunology. The authors are thankful to the DBT for a generous infrastructure grant to establish the Next Generation Sequencing facility. GCS is supported by an ICMR SRF fellowship (RMBH/FW/2020/19), UR by DBT-JRF fellowship DBT/2018/NII/1035. Some strains were provided by the CGC, which is funded by the NIH Office of Research Infrastructure Programs (P40 OD010440) and the National Bioresource Project (NBRP), Japan. The schematic representations were created with BioRender.com.

## Supplementary Information

### Supplementary Information Text

#### Supplementary Materials and Methods

##### C. elegans strain maintenance

Unless otherwise mentioned, all the *C. elegans* strains were maintained and propagated at 20°C on *E. coli* OP50 using standard procedures (1). The strains used in this study were: N2 var. Bristol: wild-type, CB1370*: daf-2(e1370)III,* GR1307: *daf-16(mgDf50) I,* HT1890: *daf-16 (mgDf50)I; daf-2 (e1370)III,* CU1546: *ced-1p::ced-1::GFP,* HT1881: *daf-16(mgDf50) I; daf-2(e1370) unc-119(ed3)III; lpIs12.lpIs12 [daf-16a::RFP + unc-119(+)],* HT1882: *daf-16(mgDf50) I; daf-2(e1370) unc-119(ed3) III; lpIs13. lpIs13 [daf-16b::CFP + unc-119(+)],* HT1883: *daf-16(mgDf50) I; daf-2(e1370) unc119(ed3)III; lpIs14. lpIs14 [daf-16f::GFP + unc-119(+)], DR1568: daf-2(e1371) III,* DR1572: *daf-2(e1368) III,* JT9609: *pdk-1(sa680)X,* TJ1052: age-1(hx546) II, KW2126: ckSi6 I; cdk-12(tm3846) III. ckSi6 [cdk-12::GFP + unc-119(+)] I, DCL569: *mkcSi13 II; rde-1(mkc36) V [mkcSi13 [sun-1p::rde-1::sun-1 3’UTR + unc-119(+)] II],* NR350: *rde-1(ne219)* V; *kzIs20[hlh-1p::rde-1 + sur-5p::NLS::GFP*], NR222: *rde-1(ne219) V; kzIs9[(pKK1260) lin-26p::NLS::GFP + (pKK1253) lin-26p::rde-1 + rol-6(su1006)], VP303: rde-1(ne219) V; kbIs7[nhx-2p::rde-1 + rol-6(su1006)], NK640: rrf-3(pk1426) II; unc-119(ed4) III; rde-1(ne219) V; qyIs102[fos-1ap::rde-1(genomic) + myo-2::YFP + unc-119(+)], TU3401: sid-1(pk3321) V; uIs69 V[pCFJ90 (myo-2p::mCherry) + unc-119p::sid-1], GR1336: daf-2(e1370) III; njEx32[ges-1p::daf-2(+) + ges-1p::GFP + rol-6(su1006)], GR1337: daf-2(e1370) III; njEx38[unc-54p::daf-2(cDNA)::unc-54 3’UTR + unc-54p::GFP + rol-6(su1006)], GR1339: daf-2(e1370) III; mgEx376[unc-14p::daf-2 + rol-6(su1006)], GR1340: daf-2(e1370) III; mgEx373[unc-119p::daf-2(cDNA)::unc-54 3’UTR + rol-6(su1006)], daf-16(mu86) I; daf-2(e1370) III, CF1515: daf-16(mu86) I;daf-2(e1370) III, Ex myo-3::daf-16, CF1514: daf-16(mu86) I;daf-2(e1370) III, Ex ges-1::daf-16, CF1442: daf-16(mu86) I;daf-2(e1370) III, Ex unc-119::daf-16, GC1285: daf-16(m26); daf-2(e1370)III, Ex fos1a::daf-16, SD551: let-60(ga89) IV, WM27: rde-1(ne219) V*. The other strains including daf-2(e1370)III; WM27: rde-1(ne219) V, daf-2(e1370)III; SD55: let-60(ga89)IV, daf-2(e1370)III; DCL569: mkcSi13 II; rde-1(mkc36) V [mkcSi13 [sun-1p::rde-1::sun-1 3’UTR + unc-119(+)] II], daf-2(e1370)III; NR350: rde-1(ne219) V; kzIs20[hlh-1p::rde-1 + sur-5p::NLS::GFP], daf-2(e1370)III; NR222: rde-1(ne219) V; kzIs9[(pKK1260) lin-26p::NLS::GFP + (pKK1253) lin-26p::rde-1 + rol-6(su1006)], daf-2(e1370)III; VP303: rde-1(ne219) V; kbIs7[nhx-2p::rde-1 + rol-6(su1006)], daf-2(e1370)III; NK640: rrf-3(pk1426) II; unc-119(ed4) III; rde-1(ne219) V; qyIs102[fos-1ap::rde-1(genomic) + myo-2::YFP + unc-119(+)], daf-2(e1370)III; TU3401: sid-1(pk3321)V; uIs69 V[pCFJ90 (myo-2p::mCherry) + unc-119p::sid-1], daf-2(e1370)III; NK640: rrf-3(pk1426) II; unc-119(ed4) III; rde-1(ne219) V; qyIs102[fos-1ap::rde-1(genomic) + myo-2::YFP + unc-119(+)] were generated in-house using standard cross-over techniques.

#### Preparation of RNAi plates

RNAi plates were poured using autoclaved nematode growth medium supplemented with 100 µg/ml ampicillin and 2 mM IPTG. Plates were dried at room temperature for 2-3 days. Bacterial culture harbouring an RNAi construct was grown in Luria Bertani (LB) media supplemented with 100 µg/ml ampicillin and 12.5 µg/ml tetracycline, overnight at 37°C in a shaker incubator. Saturated cultures were re-inoculated the next day in fresh LB media containing 100 µg/ml ampicillin by using 1/100^th^ volume of the primary inoculum and grown in 37°C shaker until OD_600_ reached 0.5-0.6. The bacterial cells were pelleted down by centrifuging the culture at 3214 g for 10 minutes at 4°C and resuspended in 1/10^th^ volume of M9 buffer containing 100 µg/ml ampicillin and 1 mM IPTG.

Strong *cdk-12* KD leads to developmental defects and the *cdk-12* mutants are non-viable. So, we diluted the *cdk-12* RNAi with control RNAi-expressing bacteria or initiated RNAi after L4, according to the experimental requirements. Different dilutions of RNAi were made by mixing with the control RNAi feed. For *cdk-12* RNAi plates, we have used 1:3 dilution of *cdk-12*:control RNAi. Around 300 µl of this suspension was seeded onto RNAi plates and left at room temperature for 2-3 days for drying, followed by storage at 4°C till further use.

#### Hypochlorite treatment to obtain eggs and synchronizing worm population

Gravid adult worms, initially grown on *E. coli* OP50 bacteria were collected using M9 buffer in a 15 ml falcon tube. Worms were washed thrice by first centrifuging at 652 g for 60 seconds followed by resuspension of the worm pellet in 1X M9 buffer. After the third wash, the worm pellet was resuspended in 3.5 ml of 1X M9 buffer and 0.5 ml 5N NaOH and 1 ml of 4% Sodium hypochlorite solution were added. The mixture was vortexed for 5-7 minutes until the entire worm bodies dissolved, leaving behind the eggs. The eggs were washed 5-6 times, by first centrifuging at 1258 g, decanting the 1X M9, followed by resuspension in fresh 1X M9 buffer to remove traces of bleach and alkali. After the final wash, eggs were kept in 15 ml falcons with ∼ 10 ml of 1X M9 buffer and kept on rotation ∼15 r.p.m for 17 hours to obtain L1 synchronized worms for all strains. The L1 worms were obtained by centrifugation at 805 g followed by resuspension in approximately 100-200 µl of M9 and added to different RNAi plates.

#### Dauer Arrest Assay

The *daf-2(e1370)* gravid adult worms, initially grown on *E. coli* OP50 were bleached and approximately 200 eggs were added to control and test RNAi plates, each in duplicates. One of the two RNAi plate containing eggs was placed at 20°C and the other at 22.5°C for each RNAi type. Animals were scored for dauer arrest when the non-dauer animals reached adulthood, 72 h or 96 h later.

#### RNA isolation

Worms grown on control or *cdk-12* RNAi were collected using 1X M9 buffer and washed thrice to remove bacteria. Trizol reagent (200 μl; Takara Bio, Kusatsu, Shiga, Japan) was added to the 50 μl worm pellet and subjected to three freeze-thaw cycles in liquid nitrogen with intermittent vortexing to break open worm bodies. The samples were then frozen in liquid nitrogen and stored at −80 °C till further use. Later, 200 μl of Trizol was again added to the worm pellet and the sample was vortexed vigorously. To this, 200 μl of chloroform was added and the tube was gently inverted several times followed by 3 minutes incubation at room temperature. The sample was then centrifuged at 12000g for 15 minutes at 4°C. The RNA containing the upper aqueous phase was gently removed into a fresh tube without disturbing the bottom layer and interphase. To this aqueous solution, an equal volume of isopropanol was added and the reaction was allowed to sit for 10 minutes at room temperature followed by centrifugation at 12000g for 10 minutes at 4°C. The supernatant was carefully discarded without disturbing the RNA-containing pellet. The pellet was washed using 1 ml 70% ethanol solution followed by centrifugation at 12000g for 5 minutes at 4°C. The RNA pellet was further dried at room temperature and later dissolved in nuclease-free water (Qiagen, Hilden, Germany) followed by incubation at 65°C for 10 minutes with intermittent tapping. The concentration of RNA was determined by measuring absorbance at 260 nm using NanoDrop UV spectrophotometer (Thermo Scientific, Waltham, USA) and quality checked using denaturing formaldehyde-agarose gel.

#### Gene expression analysis using quantitative real-time PCR (QRT-PCR)

First-strand cDNA synthesis was carried out using the Iscript cDNA synthesis kit (Biorad, Hercules, USA) following the manufacturer’s guidelines. The prepared cDNA was stored at −20°C. Gene expression levels were determined using the Brilliant III Ultra-Fast SYBR® Green QPCR master mix (Agilent, Santa Clara, USA) and Agilent AriaMx Real-Time PCR system (Agilent, Santa Clara, USA), according to manufacturer’s guidelines. The relative expression of each gene was determined by normalizing the data to actin expression levels.

#### RNAi life span

Gravid adult worms, initially grown on *E. coli* OP50 were bleached and their eggs were allowed to hatch in 1X M9 buffer for 17 hours to obtain L1 synchronized worms. The L1 worms obtained were added to control RNAi plates. On reaching adulthood, 50-60 L4 worms were transferred to the control or *cdk-12* RNAi plates in triplicates and on reaching the young adult stage, were overlaid with Fluoro-deoxyuridine (FudR) to a final concentration of 0.1 mg/ml of agar (2). On the 7^th^ Day of adulthood, sick, sluggish, and slow-dwelling worms were removed from the life span population and the remaining were considered as the number of subjects ‘N’. Following this, the number of dead worms was scored every alternate day and plotted as % survival against the number of days.

#### Heat survival assay

Worms were grown on control RNAi, L1 stage onwards, and approximately 50 L4 worms for each strain were transferred to control or *cdk-12* RNAi-seeded NGM plates in triplicates and transferred at an incubator maintained at 20 °C. About 48 hours post-transfer, the RNAi plates were transferred to an incubator maintained at 35° C. Following this survival was scored every 2 hours till all worms were dead.

#### Measurement of cell corpses using CED-1::GFP

The number of engulfed cell corpses was analyzed using CED-1::GFP expressing transgenic worms, where CED-1 is a transmembrane protein expressed on phagocytic cells that engulf cell-corpses. Transgenic worms, expressing CED-1 fused to GFP under *ced-1* promoter [*ced-1p::ced-1::GFP(smIs34)*], were bleached and their eggs were allowed to hatch in 1X M9 buffer for 17 hours to obtain L1 synchronized worms. Approximately 200 L1 worms were placed onto control or *cdk-12* RNAi in triplicates. On day 1 adult stage, worms were visualized under an LSM-980 confocal microscope (Carl Zeiss, Oberkochen, Germany). The number of cell corpses per gonad was counted.

#### DAPI staining

Worms were grown on control or *cdk-12* RNAi, L1 stage onwards. Day 1 adults were collected in 1X M9 buffer in a 1.5 ml Eppendorf tube and worms were allowed to settle down. Using a glass Pasteur pipette, the 1X M9 was discarded, leaving behind a ∼ 100 µl worm suspension. Then, 1 ml chilled 100 % methanol was added, to the worm pellet and incubated for 30 minutes at −20°C. The pellet was placed on a glass slide and Fluroshield with DAPI (Invitrogen, Carlsbad, USA) was added. For staining dissected gonads, worms were placed onto a glass slide and the gonads were obtained by cutting the pharynx or tail end of the worm using a sharp 25G needle.

After collecting gonads for 10 minutes, 500 µl chilled 100% methanol was added onto the slide and allowed to dry. Fluroshield with DAPI (Invitrogen, Carlsbad, USA) was added. The slides were imaged using a confocal microscope (Carl Zeiss, Oberkochen, Germany).

#### pMPK-1 Immunostaining staining

pMPK-1 immunostaining was performed as described previously (3). Briefly, on day 1, adult worms were dissected in 1X M9 buffer to obtain gonads. The dissections were performed on a glass slide and within a 5 minutes window to prevent the loss of pMPK-1 signal. Following this, the dissected gonads and remaining worms were transferred to a 10 ml round bottom glass tube using a glass pipette. To this, 2 ml of 3% Paraformaldehyde (PFA) was added and incubated at room temperature for 10 minutes. Next, 3 ml of 1X PBST was added for washing to remove PFA. The tube was allowed to stand till the dissected gonads and residual intact worms settled to the bottom. After removing the supernatant, the washing was repeated twice more. After the final PBST wash, 2 ml of 100% methanol was added and the tubes were incubated at −20 °C for 1 hour. Three 1X PBST washes were then given, as described previously and after the final wash, the worms were transferred to a 1.5 ml glass tube. Carefully, excess PBST was removed using a glass Pasteur pipette. Blocking was performed at room temperature for 1 hour using 100 µl of 30% Normal goat serum (NGS) per tube. After blocking, 100 µl of pMPK-1 antibody diluted (1:400) in 30% NGS was added to each tube, and tubes were capped, sealed with parafilm to prevent loss from evaporation, and stored at 4° C overnight. After three 1X PBST washes, 100 µl of secondary antibody in 30% NGS was added to each tube and incubated at room temperature for 2 hours. Again, three 1X PBST washes were given and excess PBST was removed. Glass Pasteur pipettes were used to pick the stained gonads in 1X PBST onto the glass slide. Quickly before the slides are completely dried, Fluroshield with DAPI (Invitrogen, Carlsbad, USA) was added and a coverslip was slowly placed using a needle, to avoid air gaps. The coverslip was gently pressed and edges were sealed with transparent nail paint. The slides were imaged using a confocal microscope (Carl Zeiss, Oberkochen Germany). The pMPK-1 signal was quantified using ImageJ software.

#### RNA-seq

Synchronized late-L4 worms grown on control or *cdk-12* RNAi were collected using 1X M9 buffer, after washing it thrice to remove bacteria. Total RNA was isolated from these worm pellets using the Trizol method. The concentration of RNA was determined by measuring absorbance at 260 nm using NanoDrop UV spectrophotometer (Thermo Scientific, Waltham, USA) and RNA quality was checked using RNA 6000 NanoAssay chip on a Bioanalyzer 2100 machine (Agilent Technologies, Santa Clara, USA) RNA above RNA integrity number = 8 was included for the study. In one batch the sequencing Libraries were constructed using NEBNext® Poly(A) mRNA Magnetic Isolation Module (Catalog no-E7490L, New England Biolabs, Ipswich, Massachusetts, USA) and NEBNext® Ultra™ II Directional RNA Library Prep kit (Catalog no-E7765L), according to the manufacturer’s instructions. For sequencing, equimolar quantities of all libraries were pooled and sequenced on Illumina Hiseq 2500 sequencer (Illumina Inc., San Diego, California, USA) as per manufacturer’s instructions using Hiseq Rapid v2 single end 50 cycles kit (1×50 cycles). In another independent batch, libraries were constructed using Truseq stranded mRNA library (for human/animal/plant) – and sequencing was performed in NovaSeq 6000 platform, 100bp paired-end (PE) with 30 million reads.

#### RNA-seq Analysis

Sequencing reads were subjected to quality control using the FASTQC kit. Alignment of the reads to WBcel235 genome was carried out with Tophat2 (4) version 2.1.0 with an average 95% alignment rate. No novel junctions and novel insertions-deletions were considered with the parameters “-no-novel-junc” and “no-novel-indel”, respectively. Gene counts were obtained with feature counts (5) version 1.6.3 and WBcel235 Ensembl annotation v95. Gene expression analysis was performed using DeSeq2 (6) package. Differentially expressed genes were defined as those with *P*-values below 5%. Genes with a cut-off of fold change > 2 and fold change < −2 were considered as upregulated and downregulated genes, respectively. For downstream analysis, the function variance stabilising transformations (VST) (7) in DeSeq2 package was implemented. Enrichment analysis was performed using the online tool DAVID 6.8 with a cutoff of FDR<10%. The dot plot was plotted with ggplot2 in R. The heatmap was plotted with the help of the heatmap function in R.

### List of primers used in the study

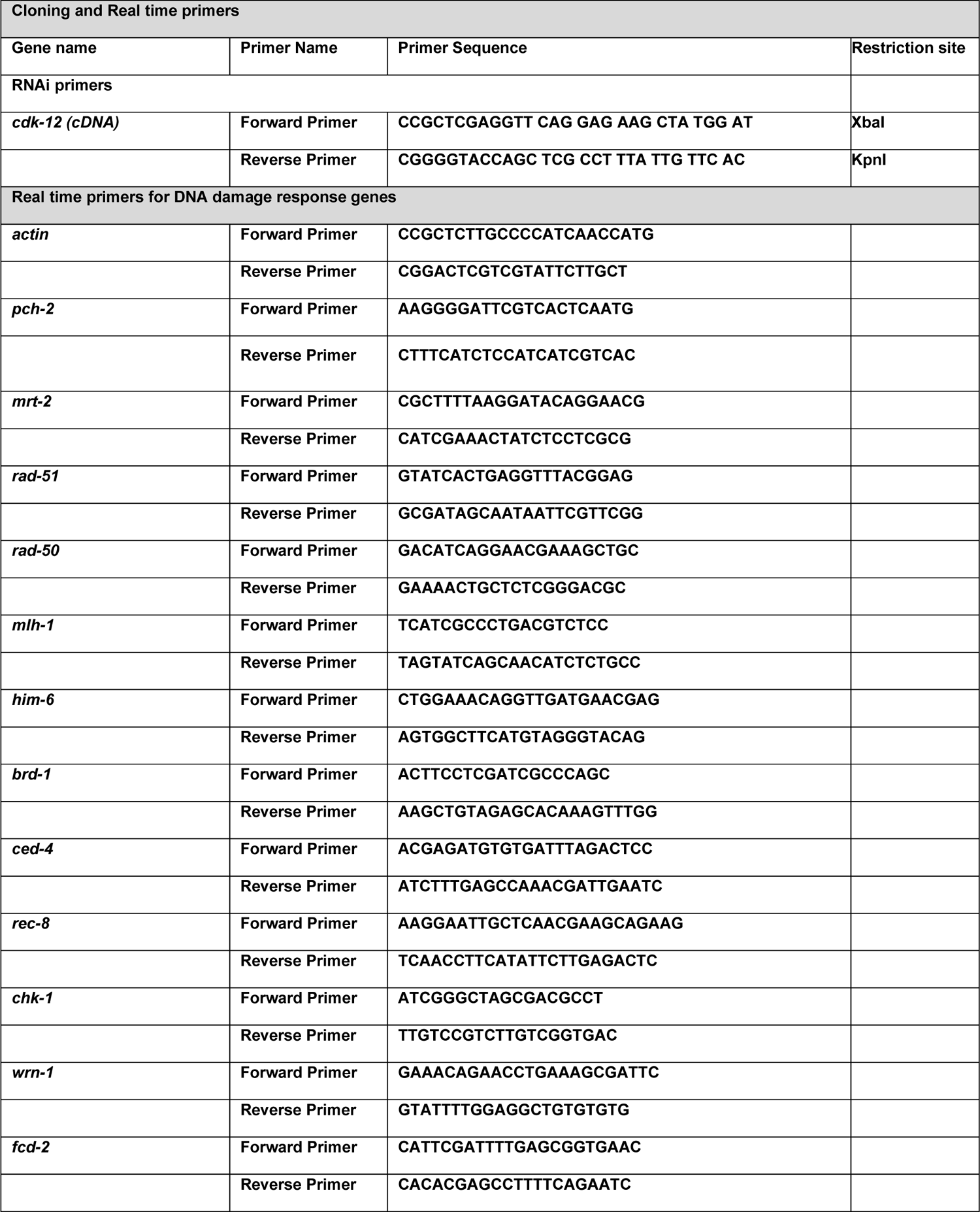
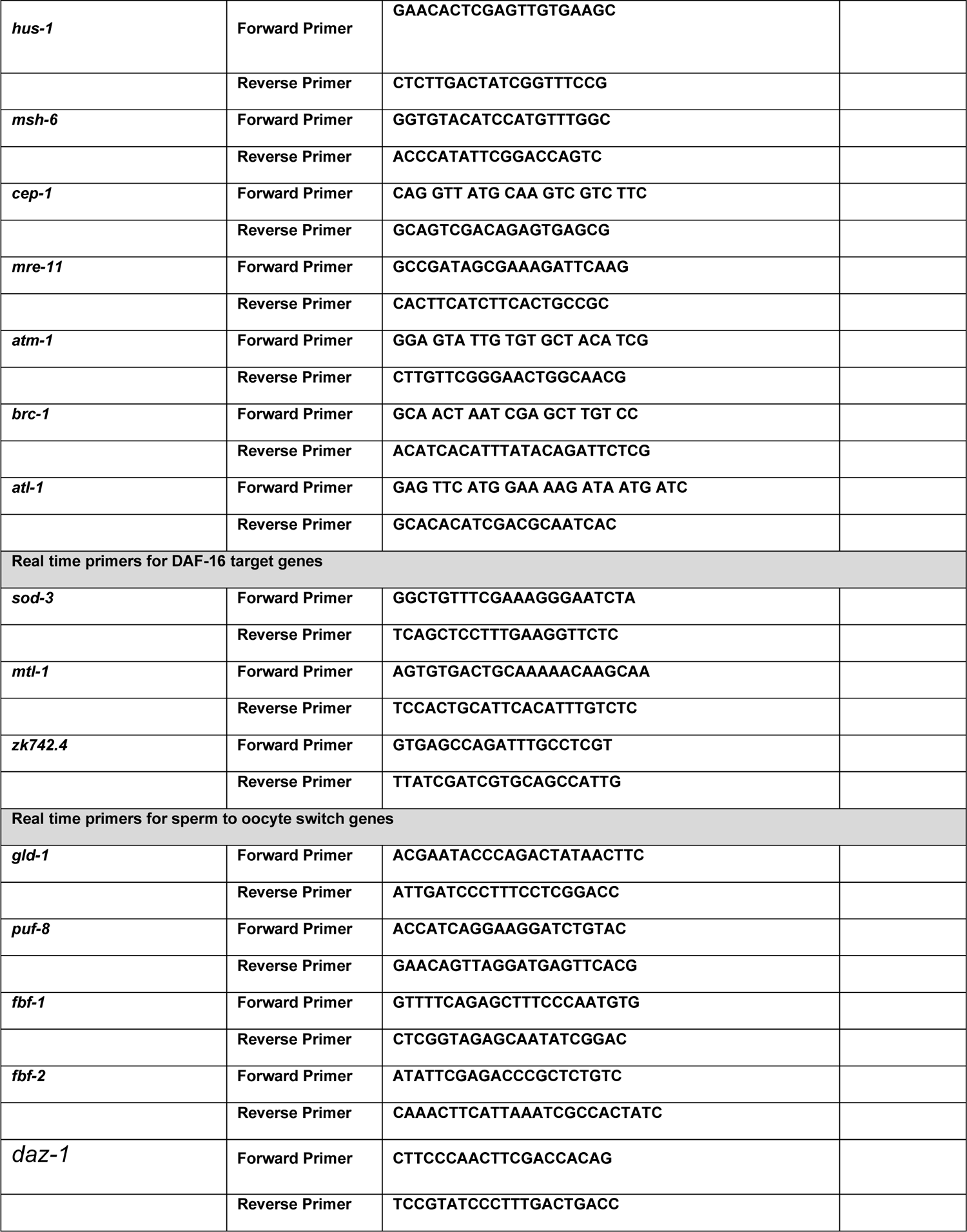
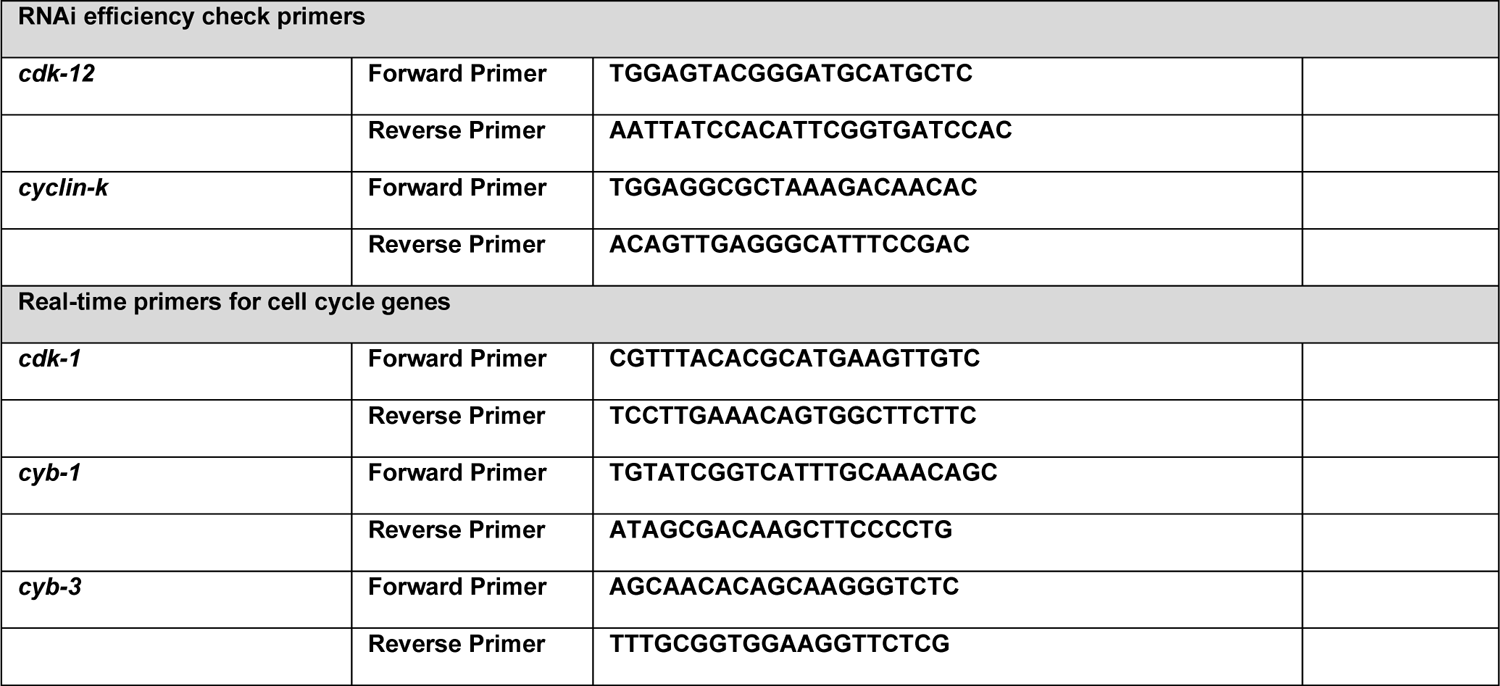

**Supplementary Figure 1.**
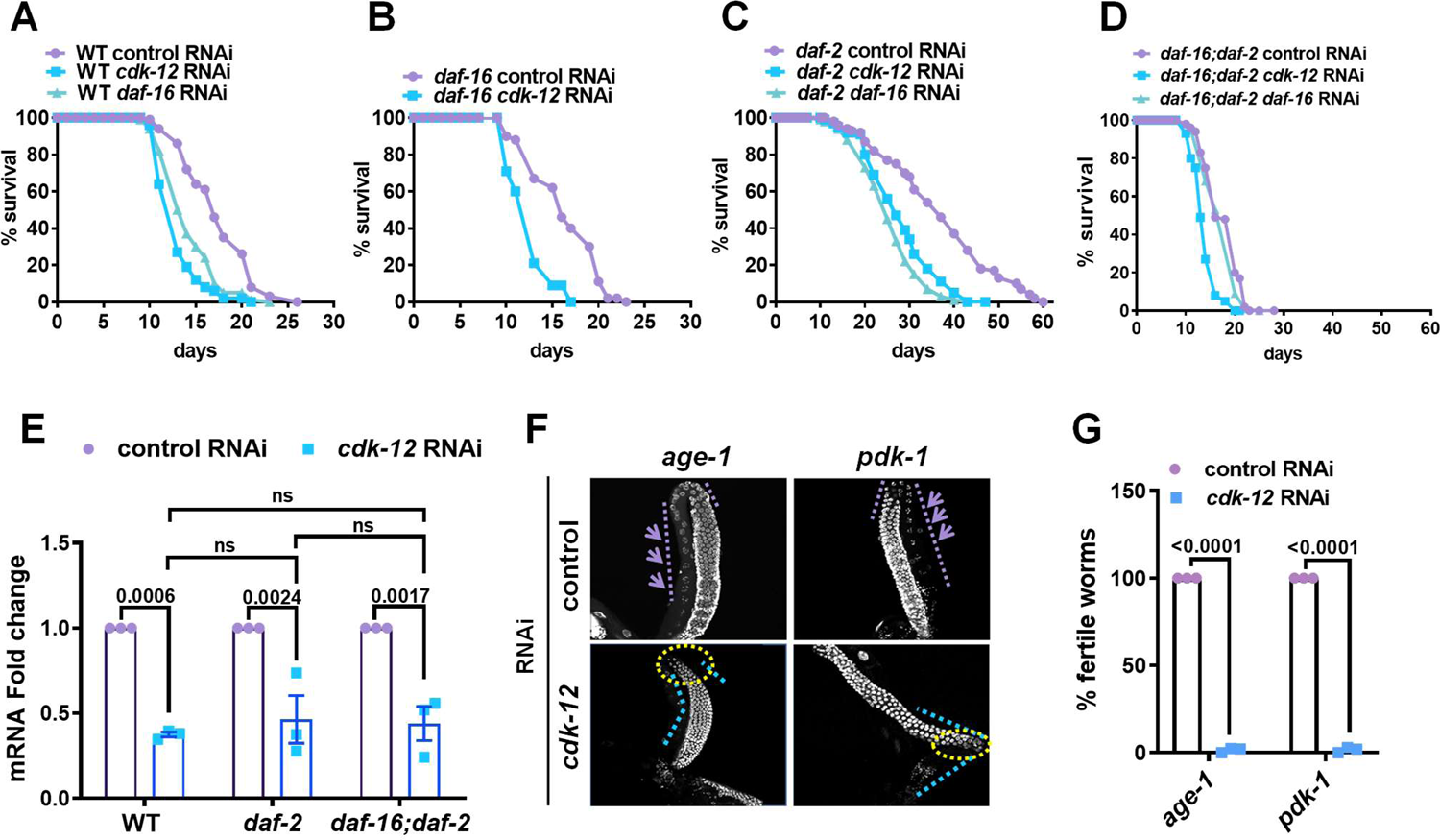
**(A-D)** The effect of *cdk-12* RNAi on lifespan of wild-type, *daf-2(e1370*), *daf-16(mgdf50)* and *daf-16(mgdf50);daf-2(e1370)*. Pooled life span from three independent biological replicates is shown. **(E)** Quantitative RT-PCR analysis showing *cdk-12* RNAi KD efficiency in WT, *daf-2(e1370)* and *daf-16(mgdf50);daf-2(e1370)*. Expression levels were normalized to *actin*. Averages of three biological replicates are shown. Two-way ANOVA-Sidak multiple comparisons test. **(F)** Representative fluorescence images of dissected gonads stained with DAPI. In *age-1(hx546)* and *pdk-1(sa680)*, germline arrests on *cdk-12* RNAi. Image were captured at 400X magnification **(G)** Percentage of fertile worms in *age-1(hx546)* and *pdk-1(sa680)* on *cdk-12* RNAi. Most worms are sterile in the two strains on *cdk-12* KD. Average of three biological replicates (n≥20 for each replicate). One way ANOVA Error bars are SEM. ns, non-significant. Experiments were performed at 20°C. Source data is provided in Dataset S1.

**Supplementary Figure 2.**
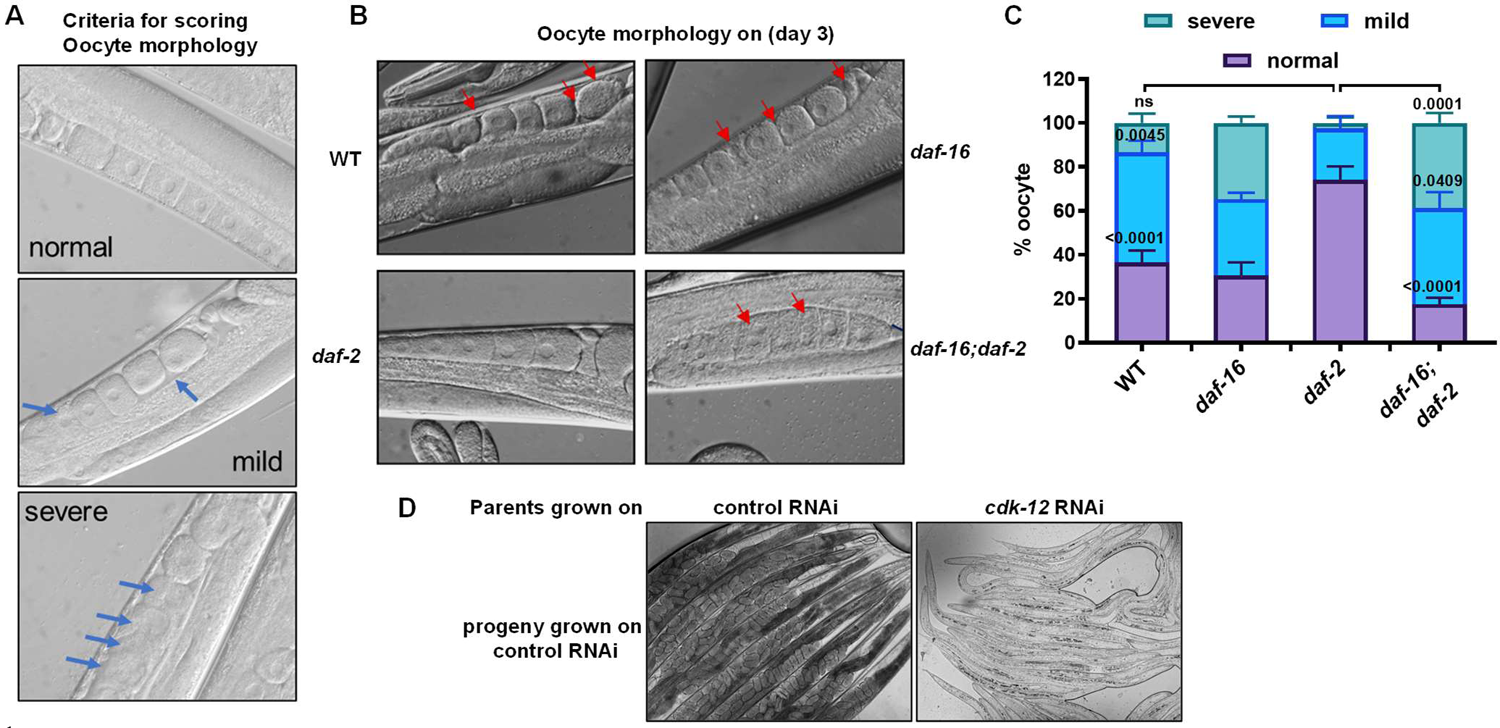
**(A)** Representative DIC images of oocyte quality that was used to set criteria for scoring in Figure S2B, S2C. Extent of deterioration of oocytes is categorized as normal, mild or severe. Arrows indicate morphologically disorganized defective oocyte. Image were captured at 400X magnification. **(B-C)** Representative images of oocyte morphology at day 3 of adulthood showing defects (B) that are quantified in (C). Strains used are WT, *daf-16(mgdf50), daf-2(e1370)* and *daf-16(mgdf50);daf-2(e1370*. Red arrows indicate defective oocytes. Average of three biological replicates (n≥30 for each replicate). Error bars are SEM. ns, non-significant. One way ANOVA. Image were captured at 400X magnification. **(D)** Representative DIC images of *daf-16(mgdf50);daf-2(e1370)* showing the developmental arrest of progeny grown on control RNAi; their parents were grown on control or *cdk-12* RNAi. Image were captured at 100X magnification. Experiments were performed at 20 °C. Source data is provided in Dataset S1.

**Supplementary Figure 3.**
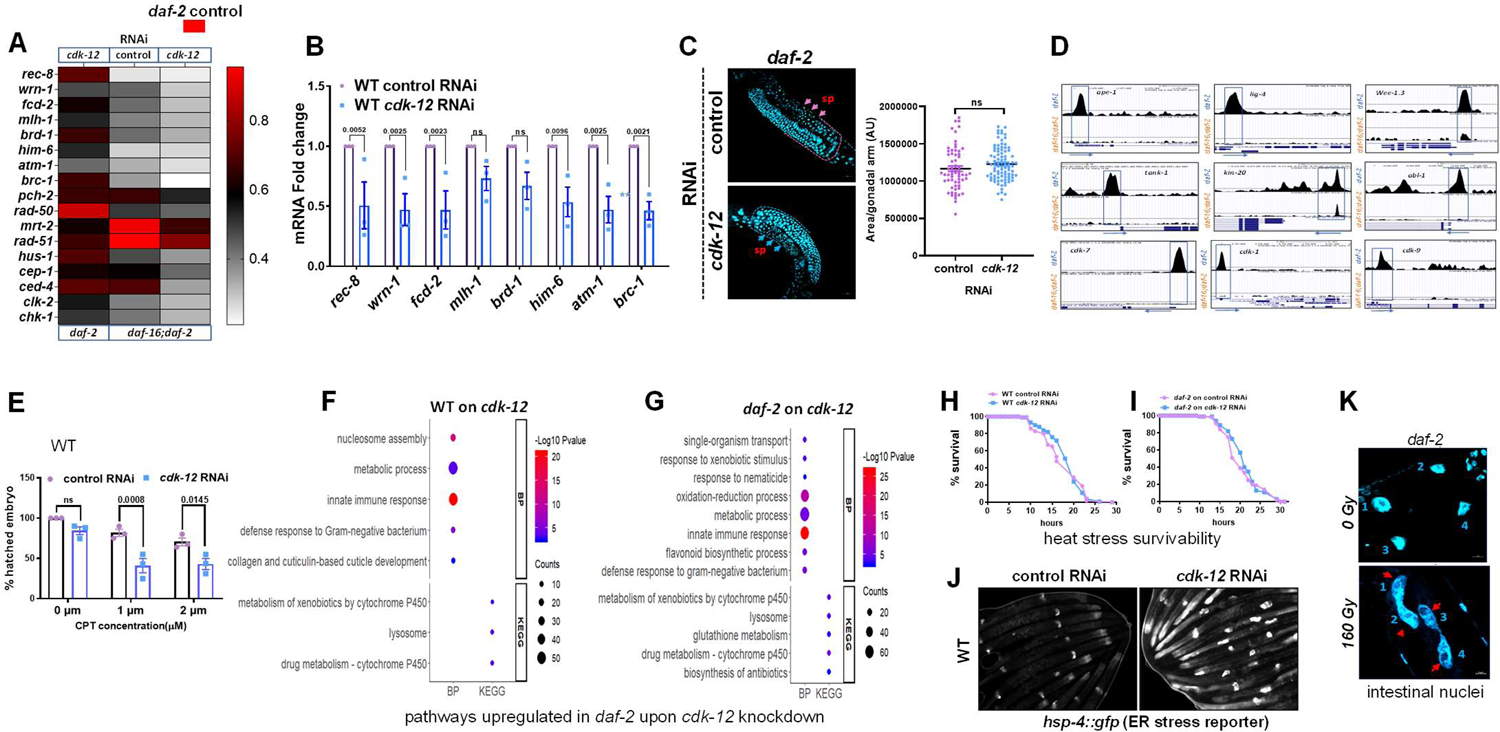
**(A)** Heat map representation of quantitative RT-PCR of DDR genes. The levels of transcripts of *daf-2(e1370)* on control RNAi was taken as 1. Levels of transcripts in *daf-2(e1370)* on *cdk-12* RNAi and *daf-16(mgdf50);daf-2(e1370)* on control or *cdk-12* RNAi are shown. Expression levels were normalized to actin. Average of three biological replicates are shown. **(B)** Quantitative RT-PCR analysis showing downregulation of DDR gene upon *cdk-12* KD in WT. Expression levels were normalized to *actin*. Averages of three or four biological replicates are shown. One way ANOVA. **(C)** Representative confocal image of DAPI stained germline at late L4 stage and its quantification in *daf-2(e1370)* on control or *cdk-12* RNAi. Germline size was quantified by measuring area (AU) using ImageJ software. Total n≈50 worms. Unpaired t test with Welch’s correction, Two-tailed. **(D)** UCSC browser view of DAF-16/FOXO peaks on promoters of DNA repair genes and cell cycle regulators as analysed by ChIP-seq of *daf-2(e1370)* and *daf-16(mgdf-50);daf-2(e1370)* strains. Blue boxes represent the promoter regions having DAF-16 binding peaks in *daf-2(e1370)*. **(E)** Decrease in the percentage of hatched embryo in WT grown on *cdk-12* RNAi upon treatment with DNA damaging agent camptothecin. Average of three biological replicates are shown (n≥20 for each replicate). One way ANOVA. **(F, G)** GO BP and KEGG pathway enrichment analysis of genes upregulated in WT (F), *daf-2(e1370)* (G), upon *cdk-12* KD, using DAVID, as compared to control RNAi. **(H, I)** Increase in heat stress (35 °C) survivability that is observed in WT (H), *daf-2(e1370)* (I) on *cdk-12* RNAi, as compared to control RNAi. Three biologically independent replicates are combined to plot the survival curves. **(J)** Representative fluorescence image showing increase in expression of *hsp-4::gfp* upon *cdk-12* KD in WT. **(K)** Representative fluorescence images of DAPI stained day-1 adult WT worms showing incomplete separation of intestinal cell nucleus upon γ-irradiation (140 Gy) at L1. Error bars are SEM. ns, non-significant. Experiments were performed at 20 °C. Source data is provided in Dataset S1.

**Supplementary 4.**
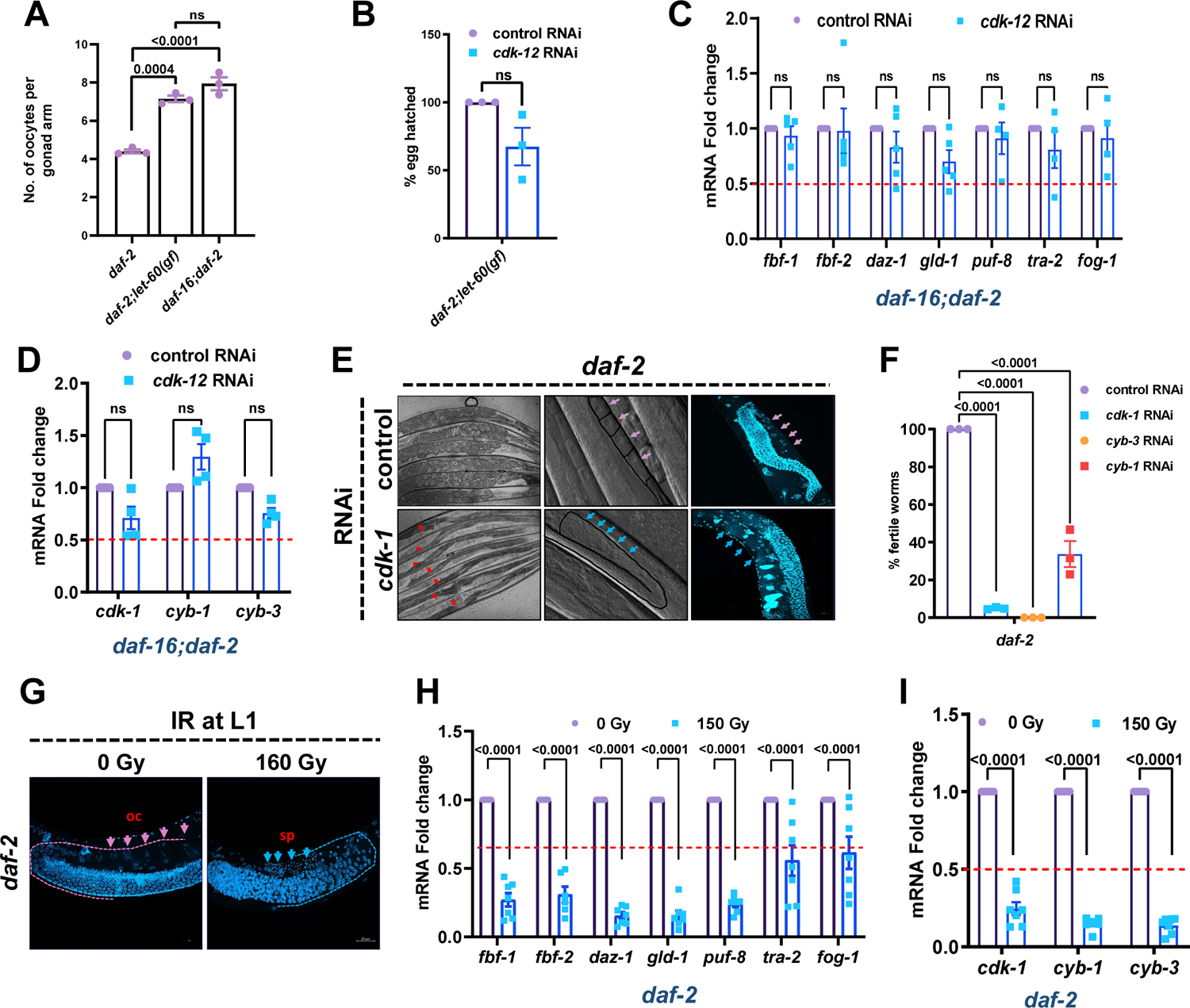
**(A)** Oocyte count per gonadal arm of *daf-2(e1370), daf-2(e1370);let-60(ga89) and daf-16(mgdf50);daf-2(e1370)*, on control RNAi. Average of three biological replicates (n≥15 for each replicate). One way ANOVA. **(B)** Percentage of eggs hatched upon *cdk-12* KD. Average of three biological replicates (n≥15 for each replicate). Unpaired t test with Welch’s correction, Two-tailed. **(C)** Quantitative RT-PCR analysis showing no significant downregulation of sperm-to-oocyte switch genes in *daf-16(mgdf50);daf-2(e1370)* upon *cdk-12* KD. Expression levels were normalized to *actin*. Average of four biological replicates are shown. One way ANOVA. **(D)** Quantitative RT-PCR analysis showing no significant downregulation of cell cycle regulator *cdk-1* and its binding partner *cyb-1* and *cyb-3* (mammalian Cyclin B orthologs) in *daf-16(mgdf50);daf-2(e1370)* upon *cdk-12* KD. Expression levels were normalized to *actin*. Average of four biological replicates are shown. One way ANOVA. **(E)** Representative images of *daf-2(e1370)* worms showing fertility, oocyte morphology, and DAPI-stained nuclei (boxed with dashed line) of the germline. The worms were grown on control and *cdk-1* RNAi. Red arrows showing sterile worms. Pink arrows showing oocytes, blue arrows point to the lack of it. **(F)** Percentage of fertile worms in *daf-2(e1370)* on control, *cdk-1*, *cyb-1* or *cyb-3* RNAi. Average of three biological replicates (n≈30 for each replicate). One way ANOVA. **(G)** Representative fluorescence images of DAPI-stained germline of day-1 adult *daf-2(e1370)* worm upon γ-irradiation (160 Gy) at L1 larval stage. Germline was arrested at pachytene stage of meiosis. Oc, oocyte (pink arrows); sp, sperms (blue arrows). **(H, I)** Quantitative RT-PCR analysis of *daf-2* worms showing significant downregulation of the sperm-to-oocyte switch genes (H) and the cell cycle regulator *cdk-1* and its binding partner *cyb-1* and *cyb-3* (I) upon γ-irradiation (160 Gy) at L1. Expression levels were normalized to *actin*. Averages of seven biological replicates are shown. Unpaired t test with Welch’s correction, Two-tailed. Error bars are SEM. ns, non-significant. Experiments were performed at 20 °C. Source data is provided in Dataset S1.

**Supplementary 5.**
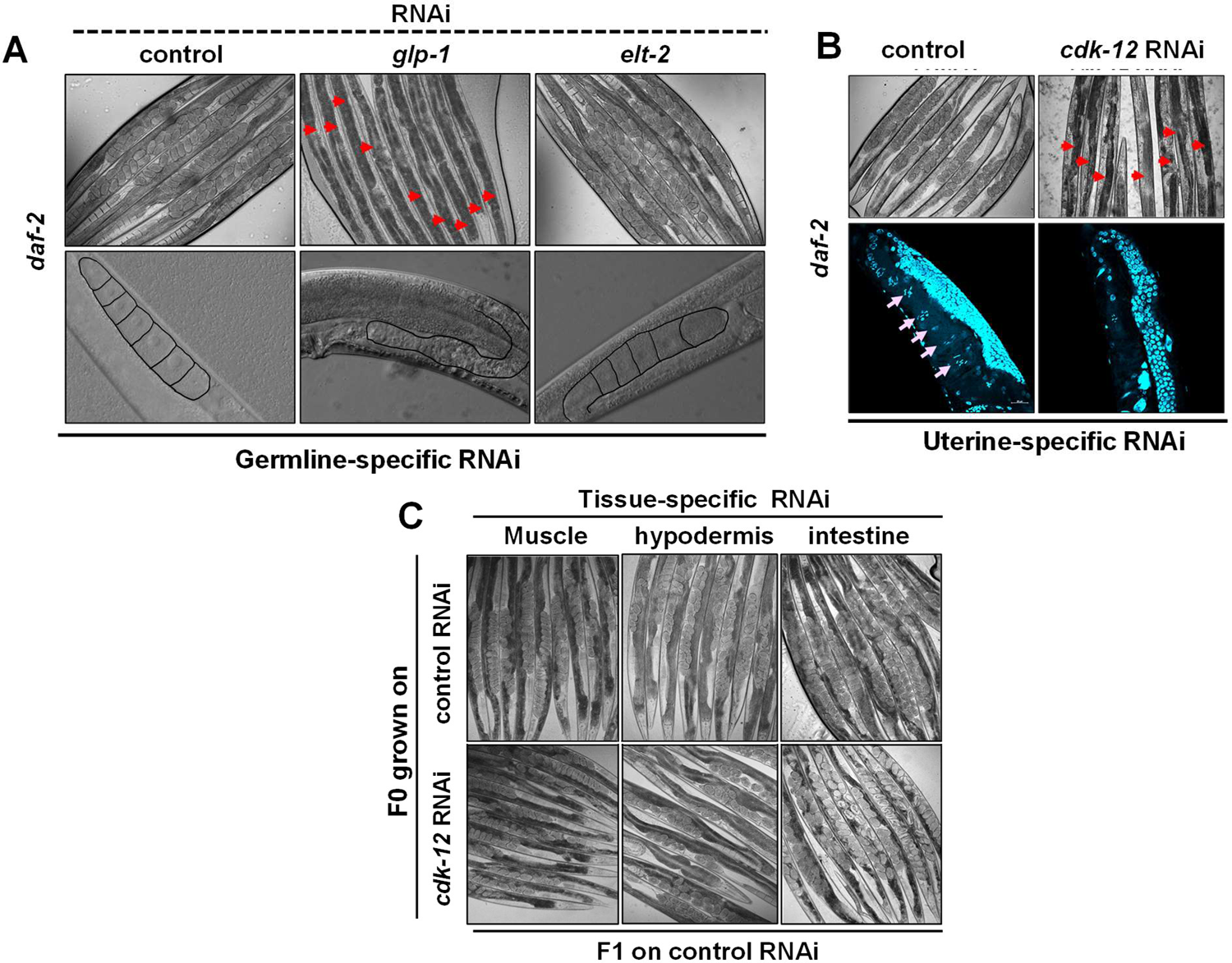
**(A)** Germline-specific KD of *elt-2* and *glp-1* in *daf-2(e1370);mkcSi13 II; rde-1(mkc36) V [mkcSi13 [sun-1p::rde-1::sun-1 3’UTR + unc-119(+)] f II]* (RNAi machinery is active only in germline). Arrows indicate sterile worms only in case of *glp-1* RNAi. Upper panels are 100x brightfield images while lower panels are 400x magnified ones. **(B)** Representative DIC (upper panel) and DAPI-stained (lower panel) images showing no oocyte formation in the *daf-2(e1370)* worms having uterine-specific knockdown of *cdk-12*. Red and pink arrows indicate sterile worms and oocytes, respectively. **(C)** Representative brightfield images showing no developmental defects in the F1 progenies grown on control RNAi. In the parental generation (F0), *cdk-12* was KD in a tissue-specific manner (intestine, hypodermis or muscle) in *daf-2(e1370)* where RNAi machinery is active only in particular tissue. Experiments were performed at 20 °C. Source data is provided in Dataset S1.

**Table S1:** Life span data used in the study

| Figure no. | Genotype | RNAi | n | Mean LS ± SEM (Days) | % Change w.r.t. control | P-value (w.r.t. control) |
| --- | --- | --- | --- | --- | --- | --- |
| Supp Figure No. 1A |  |  |  |  |  |  |
| | WT | control | 226 | 17.44 ± 0.24 | | |
| | | *cdk-12* | 258 | 13.06 ± 0.14 | -25.11 | 0.00E+00 |
| | | *daf-16* | 292 | 14.29 ± 0.16 | -18.06 | 0.00E+00 |
| Supp Figure No. 1B | *daf-16* | control | 243 | 16.39 ± 0.23 | | |
| | | *cdk-12* | 192 | 12.55 ± 0.16 | -23.42 | 0.00E+00 |
| Supp Figure No. 1C | *daf-2* | control | 410 | 36.84 ± 0.6 | | |
| | | *cdk-12* | 387 | 27.93 ± 0.39 | -24.19 | 0.00E+00 |
| | | *daf-16* | 283 | 25.59 ± 0.39 | -30.54 | 0.00E+00 |
| Supp Figure No. 1D | *daf-16;daf-2* | control | 358 | 17.61 ± 0.19 | | |
| | | *cdk-12* | 323 | 13.83 ± 0.14 | -21.46 | 0.00E+00 |
| | | *daf-16* | 144 | 17.59 ± 0.27 | -0.11 | 0.42 |

**Dataset S1.** Details of the genes that were differentially expressed in the RNA-seq analysis

**Dataset S2.** Source data file

## References

1. S. P. Jackson, J. Bartek, The DNA-damage response in human biology and disease. Nature 461, 1071–1078 (2009).

2. A. L. Lopez et al., DAF-2 and ERK couple nutrient availability to meiotic progression during Caenorhabditis elegans oogenesis. Developmental cell 27, 227–240 (2013).

3. E. J. Hubbard, D. Z. Korta, D. Dalfó, Physiological control of germline development. Advances in experimental medicine and biology 757, 101–131 (2013).

4. J. Y. Wo, A. N. Viswanathan, Impact of Radiotherapy on Fertility, Pregnancy, and Neonatal Outcomes in Female Cancer Patients. International Journal of Radiation Oncology*Biology*Physics 73, 1304–1312 (2009).

5. S. Gottlieb, G. Ruvkun, daf-2, daf-16 and daf-23: genetically interacting genes controlling Dauer formation in Caenorhabditis elegans. Genetics 137, 107–120 (1994).

6. P. L. Larsen, P. S. Albert, D. L. Riddle, Genes that regulate both development and longevity in Caenorhabditis elegans. Genetics 139, 1567–1583 (1995).

7. L. R. Baugh, P. W. Sternberg, DAF-16/FOXO regulates transcription of cki-1/Cip/Kip and repression of lin-4 during C. elegans L1 arrest. Curr Biol 16, 780–785 (2006).

8. A. Mukhopadhyay, S. W. Oh, H. A. Tissenbaum, Worming pathways to and from DAF-16/FOXO. Exp Gerontol 41, 928–934 (2006).

9. R. Y. Lee, J. Hench, G. Ruvkun, Regulation of C. elegans DAF-16 and its human ortholog FKHRL1 by the daf-2 insulin-like signaling pathway. Curr Biol 11, 1950–1957 (2001).

10. S. Ogg et al., The Fork head transcription factor DAF-16 transduces insulin-like metabolic and longevity signals in *C. elegans*. Nature 389, 994–999 (1997).

11. Z. Qin, E. J. Hubbard, Non-autonomous DAF-16/FOXO activity antagonizes age-related loss of C. elegans germline stem/progenitor cells. Nat Commun 6, 7107 (2015).

12. N. M. Templeman et al., Insulin Signaling Regulates Oocyte Quality Maintenance with Age via Cathepsin B Activity. Curr Biol 28, 753–760 e754 (2018).

13. S. Luo, G. A. Kleemann, J. M. Ashraf, W. M. Shaw, C. T. Murphy, TGF-beta and insulin signaling regulate reproductive aging via oocyte and germline quality maintenance. Cell 143, 299–312 (2010).

14. C. A. Wolkow, K. D. Kimura, M. S. Lee, G. Ruvkun, Regulation of C. elegans life-span by insulinlike signaling in the nervous system. Science 290, 147–150 (2000).

15. K. Lin, H. Hsin, N. Libina, C. Kenyon, Regulation of the Caenorhabditis elegans longevity protein DAF-16 by insulin/IGF-1 and germline signaling. Nat Genet 28, 139–145 (2001).

16. B. Bartkowiak et al., CDK12 is a transcription elongation-associated CTD kinase, the metazoan ortholog of yeast Ctk1. Genes Dev 24, 2303–2316 (2010).

17. S. J. Dubbury, P. L. Boutz, P. A. Sharp, CDK12 regulates DNA repair genes by suppressing intronic polyadenylation. Nature 564, 141–145 (2018).

18. J. Austin, J. Kimble, glp-1 is required in the germ line for regulation of the decision between mitosis and meiosis in C. elegans. Cell 51, 589–599 (1987).

19. B. Kraemer et al., NANOS-3 and FBF proteins physically interact to control the sperm-oocyte switch in Caenorhabditis elegans. Curr Biol 9, 1009–1018 (1999).

20. J. Kimble, S. L. Crittenden, Controls of germline stem cells, entry into meiosis, and the sperm/oocyte decision in Caenorhabditis elegans. Annu Rev Cell Dev Biol 23, 405–433 (2007).

21. J. Z. Morris, H. A. Tissenbaum, G. Ruvkun, A phosphatidylinositol-3-OH kinase family member regulating longevity and diapause in *Caenorhabditis elegans*. Nature 382, 536–539 (1996).

22. S. Paradis, M. Ailion, A. Toker, J. H. Thomas, G. Ruvkun, A PDK1 homolog is necessary and sufficient to transduce AGE-1 PI3 kinase signals that regulate diapause in Caenorhabditis elegans. Genes Dev 13, 1438–1452 (1999).

23. E. S. Kwon, S. D. Narasimhan, K. Yen, H. A. Tissenbaum, A new DAF-16 isoform regulates longevity. Nature 466, 498–502 (2010).

24. S. T. Henderson, T. E. Johnson, daf-16 integrates developmental and environmental inputs to mediate aging in the nematode Caenorhabditis elegans. Curr Biol 11, 1975–1980 (2001).

25. N. Libina, J. R. Berman, C. Kenyon, Tissue-specific activities of C. elegans DAF-16 in the regulation of lifespan. Cell 115, 489–502 (2003).

26. K. Iwasaki, J. McCarter, R. Francis, T. Schedl, emo-1, a Caenorhabditis elegans Sec61p gamma homologue, is required for oocyte development and ovulation. J Cell Biol 134, 699–714 (1996).

27. K. M. Ekumi et al., Ovarian carcinoma CDK12 mutations misregulate expression of DNA repair genes via deficient formation and function of the Cdk12/CycK complex. Nucleic Acids Res 43, 2575–2589 (2015).

28. P. M. Joshi, S. L. Sutor, C. J. Huntoon, L. M. Karnitz, Ovarian cancer-associated mutations disable catalytic activity of CDK12, a kinase that promotes homologous recombination repair and resistance to cisplatin and poly(ADP-ribose) polymerase inhibitors. J Biol Chem 289, 9247–9253 (2014).

29. I. Clejan, J. Boerckel, S. Ahmed, Developmental modulation of nonhomologous end joining in Caenorhabditis elegans. Genetics 173, 1301–1317 (2006).

30. M. A. Ermolaeva et al., DNA damage in germ cells induces an innate immune response that triggers systemic stress resistance. Nature 501, 416–420 (2013).

31. J. Deng, X. Bai, H. Tang, S. Pang, DNA damage promotes ER stress resistance through elevation of unsaturated phosphatidylcholine in Caenorhabditis elegans. J Biol Chem 296, 100095 (2021).

32. E. M. Hedgecock, J. G. White, Polyploid tissues in the nematode Caenorhabditis elegans. Dev Biol 107, 128–133 (1985).

33. M. Kniazeva, G. Ruvkun, Rhizobium induces DNA damage in Caenorhabditis elegans intestinal cells. Proc Natl Acad Sci U S A 116, 3784–3792 (2019).

34. A. P. Bailly et al., The Caenorhabditis elegans homolog of Gen1/Yen1 resolvases links DNA damage signaling to DNA double-strand break repair. PLoS Genet 6, e1001025 (2010).

35. M. H. Lee et al., Multiple functions and dynamic activation of MPK-1 extracellular signal-regulated kinase signaling in Caenorhabditis elegans germline development. Genetics 177, 2039–2062 (2007).

36. A. L. Lopez 3rd, et al., DAF-2 and ERK couple nutrient availability to meiotic progression during Caenorhabditis elegans oogenesis. Dev Cell 27, 227-240 (2013).

37. D. L. Church, K. L. Guan, E. J. Lambie, Three genes of the MAP kinase cascade, mek-2, mpk-1/sur-1 and let-60 ras, are required for meiotic cell cycle progression in Caenorhabditis elegans. Development 121, 2525–2535 (1995).

38. D. S. Yoon, M. A. Alfhili, K. Friend, M. H. Lee, MPK-1/ERK regulatory network controls the number of sperm by regulating timing of sperm-oocyte switch in C. elegans germline. Biochemical and biophysical research communications 491, 1077–1082 (2017).

39. M. van der Voet, M. A. Lorson, D. G. Srinivasan, K. L. Bennett, S. van den Heuvel, C. elegans mitotic cyclins have distinct as well as overlapping functions in chromosome segregation. Cell cycle 8, 4091–4102 (2009).

40. L. Zou et al., Construction of a germline-specific RNAi tool in C. elegans. Sci Rep 9, 2354 (2019).

41. A. S. Pepper, D. J. Killian, E. J. Hubbard, Genetic analysis of Caenorhabditis elegans glp-1 mutants suggests receptor interaction or competition. Genetics 163, 115–132 (2003).

42. F. G. Mann, E. L. Van Nostrand, A. E. Friedland, X. Liu, S. K. Kim, Deactivation of the GATA Transcription Factor ELT-2 Is a Major Driver of Normal Aging in C. elegans. PLoS Genet 12, e1005956 (2016).

43. H. Qadota et al., Establishment of a tissue-specific RNAi system in C. elegans. Gene 400, 166–173 (2007).

44. M. V. Espelt, A. Y. Estevez, X. Yin, K. Strange, Oscillatory Ca2+ signaling in the isolated Caenorhabditis elegans intestine: role of the inositol-1,4,5-trisphosphate receptor and phospholipases C beta and gamma. J Gen Physiol 126, 379-392 (2005).

45. A. Calixto, D. Chelur, I. Topalidou, X. Chen, M. Chalfie, Enhanced neuronal RNAi in C. elegans using SID-1. Nat Methods 7, 554–559 (2010).

46. T. N. Medwig-Kinney et al., A developmental gene regulatory network for C. elegans anchor cell invasion. Development 147 (2020).

47. K. D. Kimura, D. L. Riddle, G. Ruvkun, The C. elegans DAF-2 insulin-like receptor is abundantly expressed in the nervous system and regulated by nutritional status. Cold Spring Harb Symp Quant Biol 76, 113–120 (2011).

48. D. Blazek et al., The Cyclin K/Cdk12 complex maintains genomic stability via regulation of expression of DNA damage response genes. Genes Dev 25, 2158–2172 (2011).

49. D. Blazek, The cyclin K/Cdk12 complex: an emerging new player in the maintenance of genome stability. Cell cycle 11, 1049–1050 (2012).

50. J. C. Harrison, J. E. Haber, Surviving the breakup: the DNA damage checkpoint. Annu Rev Genet 40, 209–235 (2006).

51. J. E. Cleaver, E. T. Lam, I. Revet, Disorders of nucleotide excision repair: the genetic and molecular basis of heterogeneity. Nat Rev Genet 10, 756–768 (2009).

52. I. Adriaens, J. Smitz, P. Jacquet, The current knowledge on radiosensitivity of ovarian follicle development stages. Hum Reprod Update 15, 359–377 (2009).

53. H. Tran et al., DNA repair pathway stimulated by the forkhead transcription factor FOXO3a through the Gadd45 protein. Science 296, 530–534 (2002).

54. J. Y. Yang et al., ERK promotes tumorigenesis by inhibiting FOXO3a via MDM2-mediated degradation. Nature cell biology 10, 138–148 (2008).

55. H. A. Miller, E. S. Dean, S. D. Pletcher, S. F. Leiser, Cell non-autonomous regulation of health and longevity. Elife 9 (2020).

56. H. Hsin, C. Kenyon, Signals from the reproductive system regulate the lifespan of C. elegans. Nature 399, 362–366 (1999).

57. N. Arantes-Oliveira, J. Apfeld, A. Dillin, C. Kenyon, Regulation of life-span by germ-line stem cells in Caenorhabditis elegans. Science 295, 502–505 (2002).

58. M. Levi-Ferber et al., It’s all in your mind: determining germ cell fate by neuronal IRE-1 in C. elegans. PLoS Genet 10, e1004747 (2014).

## SI References

1. T. Stiernagle, Maintenance of C. elegans. WormBook 10.1895/wormbook.1.101.1, 1-11 (2006).

2. R. Hosono, Y. Mitsui, Y. Sato, S. Aizawa, J. Miwa, Life span of the wild and mutant nematode Caenorhabditis elegans. Effects of sex, sterilization, and temperature. Exp Gerontol 17, 163–172 (1982).

3. A. L. Gervaise, S. Arur, Spatial and Temporal Analysis of Active ERK in the C. elegans Germline. J Vis Exp 10.3791/54901 (2016).

4. D. Kim et al., TopHat2: accurate alignment of transcriptomes in the presence of insertions, deletions and gene fusions. Genome Biol 14, R36 (2013).

5. Y. Liao, G. K. Smyth, W. Shi, featureCounts: an efficient general purpose program for assigning sequence reads to genomic features. Bioinformatics 30, 923–930 (2014).

6. M. I. Love, W. Huber, S. Anders, Moderated estimation of fold change and dispersion for RNA-seq data with DESeq2. Genome Biol 15, 550 (2014).

7. S. Anders, W. Huber, Differential expression analysis for sequence count data. Genome Biol 11, R106 (2010).

